# Human cortical expansion involves diversification and specialization of supragranular intratelencephalic-projecting neurons

**DOI:** 10.1101/2020.03.31.018820

**Authors:** Jim Berg, Staci A. Sorensen, Jonathan T. Ting, Jeremy A. Miller, Thomas Chartrand, Anatoly Buchin, Trygve E. Bakken, Agata Budzillo, Nick Dee, Song-Lin Ding, Nathan W. Gouwens, Rebecca D. Hodge, Brian Kalmbach, Changkyu Lee, Brian R. Lee, Lauren Alfiler, Katherine Baker, Eliza Barkan, Allison Beller, Kyla Berry, Darren Bertagnolli, Kris Bickley, Jasmine Bomben, Thomas Braun, Krissy Brouner, Tamara Casper, Peter Chong, Kirsten Crichton, Rachel Dalley, Rebecca de Frates, Tsega Desta, Samuel Dingman Lee, Florence D’Orazi, Nadezhda Dotson, Tom Egdorf, Rachel Enstrom, Colin Farrell, David Feng, Olivia Fong, Szabina Furdan, Anna A. Galakhova, Clare Gamlin, Amanda Gary, Alexandra Glandon, Jeff Goldy, Melissa Gorham, Natalia A. Goriounova, Sergey Gratiy, Lucas Graybuck, Hong Gu, Kristen Hadley, Nathan Hansen, Tim S. Heistek, Alex M. Henry, Djai B. Heyer, DiJon Hill, Chris Hill, Madie Hupp, Tim Jarsky, Sara Kebede, Lisa Keene, Lisa Kim, Mean-Hwan Kim, Matthew Kroll, Caitlin Latimer, Boaz P. Levi, Katherine E. Link, Matthew Mallory, Rusty Mann, Desiree Marshall, Michelle Maxwell, Medea McGraw, Delissa McMillen, Erica Melief, Eline J. Mertens, Leona Mezei, Norbert Mihut, Stephanie Mok, Gabor Molnar, Alice Mukora, Lindsay Ng, Kiet Ngo, Philip R. Nicovich, Julie Nyhus, Gaspar Olah, Aaron Oldre, Victoria Omstead, Attila Ozsvar, Daniel Park, Hanchuan Peng, Trangthanh Pham, Christina A. Pom, Lydia Potekhina, Ramkumar Rajanbabu, Shea Ransford, David Reid, Christine Rimorin, Augustin Ruiz, David Sandman, Josef Sulc, Susan M. Sunkin, Aaron Szafer, Viktor Szemenyei, Elliot R. Thomsen, Michael Tieu, Amy Torkelson, Jessica Trinh, Herman Tung, Wayne Wakeman, Katelyn Ward, René Wilbers, Grace Williams, Zizhen Yao, Jae-Geun Yoon, Costas Anastassiou, Anton Arkhipov, Pal Barzo, Amy Bernard, Charles Cobbs, Philip C. de Witt Hamer, Richard G. Ellenbogen, Luke Esposito, Manuel Ferreira, Ryder P. Gwinn, Michael J. Hawrylycz, Patrick R. Hof, Sander Idema, Allan R. Jones, C.Dirk Keene, Andrew L. Ko, Gabe J. Murphy, Lydia Ng, Jeffrey G. Ojemann, Anoop P. Patel, John W. Phillips, Daniel L. Silbergeld, Kimberly Smith, Bosiljka Tasic, Rafael Yuste, Idan Segev, Christiaan P.J. de Kock, Huibert D. Mansvelder, Gabor Tamas, Hongkui Zeng, Christof Koch, Ed S. Lein

**Author notes:** These authors contributed equally (Jim Berg, Staci A. Sorensen, Jonathan T. Ting, Jeremy A. Miller).

## Abstract

The neocortex is disproportionately expanded in human compared to mouse, both in its total volume relative to subcortical structures and in the proportion occupied by supragranular layers that selectively make connections within the cortex and other telencephalic structures. Single-cell transcriptomic analyses of human and mouse cortex show an increased diversity of glutamatergic neuron types in supragranular cortex in human and pronounced gradients as a function of cortical depth. To probe the functional and anatomical correlates of this transcriptomic diversity, we describe a robust Patch-seq platform using neurosurgically-resected human tissues. We characterize the morphological and physiological properties of five transcriptomically defined human glutamatergic supragranular neuron types. Three of these types have properties that are specialized compared to the more homogeneous properties of transcriptomically defined homologous mouse neuron types. The two remaining supragranular neuron types, located exclusively in deep layer 3, do not have clear mouse homologues in supragranular cortex but are transcriptionally most similar to deep layer mouse intratelencephalic-projecting neuron types. Furthermore, we reveal the transcriptomic types in deep layer 3 that express high levels of non-phosphorylated heavy chain neurofilament protein that label long-range neurons known to be selectively depleted in Alzheimer’s disease. Together, these results demonstrate the power of transcriptomic cell type classification, provide a mechanistic underpinning for increased complexity of cortical function in human cortical evolution, and implicate discrete transcriptomic cell types as selectively vulnerable in disease.

## Introduction

The neocortex is responsible for many aspects of cognitive function and is affected in numerous neurological and neuropsychiatric diseases. Great progress has been made in understanding the cell types that make up functional cortical circuitry in rodents ^1,2,3^, but our understanding of cortical cell types in human is far more rudimentary due to the relative inaccessibility of human brain tissues. A striking feature of the neocortex is its disproportionate expansion in surface area, volume, and neuron number in large-brain mammals when compared to the expansion measured in subcortical structures ^4,5^. In addition, the basic cortical architecture in primates, including human, shows a disproportionate increase in the upper or supragranular layers ^6^, whose glutamatergic (excitatory pyramidal projection) neurons make connections to other cortical and telencephalic brain regions.

The supragranular cortex in human has been historically divided into layer 2 (L2) and 3 (with further subdivision of L3 depending on the cortical area), whereas such distinctions are not possible in mouse cortex, where supragranular cortex is referred to as layer 2/3 (L2/3). At the cellular level, rodent L2/3 pyramidal neurons form a relatively homogeneous population based on electrophysiological and morphological properties ^1,2,7^, whereas in primates there is clear heterogeneity of neuron density, size, morphology, electrophysiology, and gene expression as a function of cortical depth and projection target ^8,9,10,11,12,4,13,14,15^. For example two main anatomical types have been described in human that differ in their dendritic morphology (slender-versus profuse-tufted ^13^). Many intrinsic electrophysiological properties show striking variation as a function of depth in supragranular cortex, including h-channel function that may facilitate faithful transmission of signals for neurons with long apical dendrites ^12^. Finally, very large neurons in deeper L3 of non-human primates send long-range (especially ipsilateral) corticocortical projections and express the non-phosphorylated form of heavy chain neurofilament protein, as they are immunoreactive to antibody SMI-32 (SMI-32ir) ^16^. This SMI-32ir neuron population is preferentially vulnerable to early degeneration and dramatically reduced in late-stage Alzheimer’s disease ^17,18^. Together, these observations suggest that the expansion of supragranular cortex in primate evolution supports increased complexity of corticocortical circuits, and some of these neuron types show a differential vulnerability in human neurodegenerative diseases.

Single-cell and single-nucleus RNA sequencing (RNA-seq) provides a novel technological and conceptual approach to analyze neuronal diversity and to directly target the expanded supragranular layers at the level of circuit components ^19,20,21^. Recent studies using these methods provide a comprehensive taxonomy of cell types in mouse and human cortex ^21,19^ and allow the quantitative alignment of cell types across species based on conserved gene expression. Of the approximately 100 transcriptomically-defined cell types (t-types) described per cortical structure in mouse cortex, three glutamatergic neuron t-types were found in L2/3 ^22^. Human L2 and L3 were similarly composed largely of three abundant glutamatergic t-types, with one of the main types exhibiting striking variation as a function of cortical depth ^11^. Alignment of these transcriptomic cell types between species showed that all human supragranular glutamatergic neuron types mapped to intratelencephalic (IT) projection neuron types in mouse, with the three most abundant human and mouse types all mapping to a single type in cross-species alignment ^11^. In addition to these matched types, which we refer to as “homologous” types, several glutamatergic neuron types were observed in deep L3 of human cortex that were not found in mouse supragranular cortex. Two of these t-types were most like IT neuron types located in mouse L5 and L6. Three additional human t-types found in L3-5 mapped best to mouse L4 t-types and likely represent the diffuse boundary between L3 and L4 in human cortex. The increased transcriptomic diversity of glutamatergic IT types compared to rodent suggests that human supragranular cortex may have other divergent cellular properties.

To test whether these transcriptomically defined cell types represent functional and anatomical differentiation between species, we developed a robust technology platform to apply the Patch-seq method ^23,24,25^ to human cortical tissues from neurosurgical resections and directly characterized the physiological and morphological properties of supragranular neurons. We demonstrate that the transcriptomic classification is highly correlated with other features of human glutamatergic neurons, both for different neuron types and for variation as a function of cortical depth within type. Homologous supragranular glutamatergic neuron types are more phenotypically diversified, or specialized, from one another in human compared to mouse. The most abundant neuron type shows graded characteristics in transcriptomic, physiological and morphological properties as a function of cortical depth. Finally, increased supragranular glutamatergic neuron cell type diversity is seen in human cortex with the addition of distinctive neuron types in deep L3 that correspond with the vulnerable neuron populations described in Alzheimer’s disease.

## Results

### Human supragranular cortex is more diverse than in mouse

The expansion of supragranular layers of the cortex in human compared to mouse ^4,26^ is also accompanied by major differences in cell density and neuron size. Here we compare human middle temporal gyrus (MTG) and mouse primary visual cortex (VISp). We chose this mouse region because of its rich transcriptomic characterization ^19,22^. Although we would prefer to compare identical regions, the magnitude of differential gene expression between mouse regions is far less than the difference seen between species ^11^ thus these data facilitate an informative cross-species comparison. We first characterized supragranular layers of cortex based on histology. The combined thickness of the relatively thin L2 and very thick L3 in human cortex (1.23 ± 0.15 mm) is on average 1.16 times greater than the thickness of the entire mouse cortex (1.06 ± 0.01 mm; Fig. 1a). Using neuronal (NeuN+) labeling in 25 μm sections, the average density of human neurons was 27.7 ± 4.5 thousand cells/mm^3^ (Fig. 1b; left). This distribution is not homogeneous, with higher density in L2 that decreases by half to reach a low point in mid L3 (Fig. 1b; right). In contrast, supragranular layers of mouse VISp show 6 times the neuronal density of human (165 ± 24.9 thousand cells/mm^3^) with a homogeneous distribution across cortical depth. These results are generally consistent with reported values and distributions for mouse and human ^4^. L3 is often divided into 3A, B, and C based on cytoarchitecture ^27^, but we did not observe sharp changes in cell density or soma size that demarcate subdivisions of L3. Instead, the average cross-sectional area of neuron somata doubles from L2 to deep L3 in human supragranular cortex in a graded fashion but in mouse is remarkably uniform across the depth of supraganular cortex (Fig. 1c, left). The interquartile range of mouse and human somata were 44 - 96μm^2^ and 107 - 253μm^2^, respectively, with the largest human somata exceeding 850μm^2^. Furthermore, variation in deep L3 neuron soma size is four-fold higher in human compared to mouse (Fig. 1c, right); this is clearly visible in human histological sections with large and small neurons co-mingling (Fig. 1a). Although NeuN does not distinguish glutamatergic pyramidal neurons from GABAergic inhibitory neurons, the largest neurons in human supragranular layers are pyramidal in shape (Fig. 1a).

**Figure 1:**
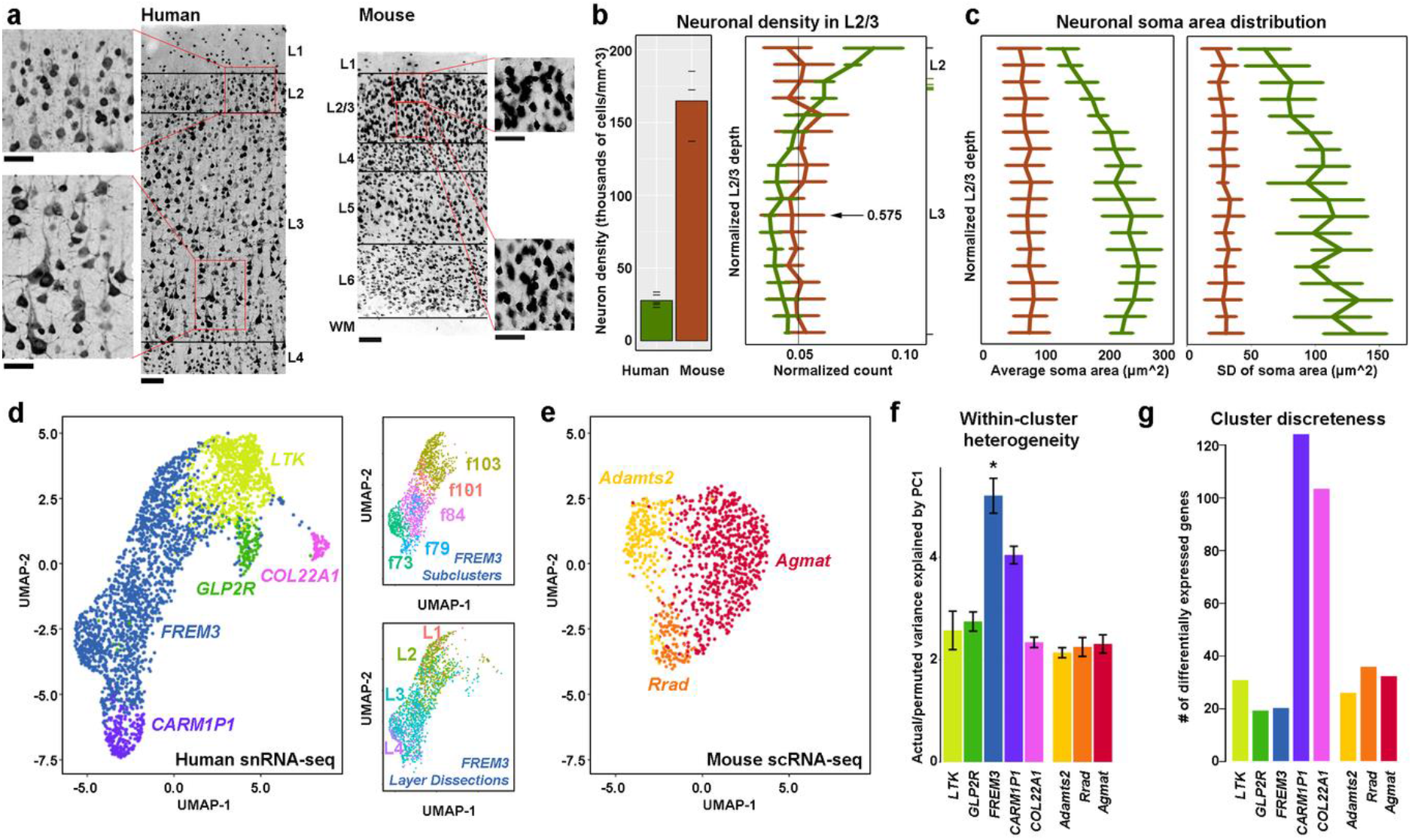
Comparison of human versus mouse supragranular neurons. **a**) NeuN IHC labeling of neurons in human MTG (left, layers 1-4) and mouse VISp (right, all layers). Higher magnification insets in upper L2 and deep L3 illustrate the much larger soma size and variability in human compared to mouse, particularly in L3. Scale bar: 100μm, main panels; 50μm insets. **b**) Left panel: human neuron density through L2 and L3 is much lower than mouse. Tick marks show individual donors. Right panel: Normalized histogram of neuron density in mouse (red) and human (green) L2/3. The minimum density in human (arrow) separates superficial and deep L3. **c**) Mean (left panel) and standard deviation (right panel) of soma area are uniform throughout the depth of mouse L2/3 but increase with depth in human. Normalized L2/3 depth is defined as the distance from the L1 - L2 (or layer 2/3 in mouse) border to the soma divided by the total thickness of L2 and L3 combined. Green tick marks on right Y axis indicate border between L2 and 3 for each human case analyzed. Error bars in **b** and **c** are SD of metrics across donors. **d**) UMAP of 2,948 dissociated human nuclei collected ^11^ from L2 and L3 of human MTG using the top 2,000 most binary genes by beta score. Cells are color-coded by t-type, with only cells mapping to the five L2 and L3 glutamatergic types included. Insets show relevant FREM3 nuclei, color coded either by subtype assignment ^11^ or by dissected layer. Note that not all FREM3 cells are assigned to a subtype. **e**) Comparable UMAP of 981 mouse cells ^22^ mapping to the three glutamatergic L2/3 neuron types in VISp. **f**) Human FREM3 t-type shows significantly more within-type heterogeneity than any other human or mouse t-type. Bar plots show average variance explained by PC1 across 100 subsets of actual versus permuted data (see Methods). Error bars show SD. **g**) Average number of DE genes between the indicated clusters and all other homologous human or mouse t-types. CARM1P1 and COL22A1 have more DE genes than other human or mouse types.

There is a similarly higher diversity of molecularly-defined, glutamatergic t-types present in human compared to mouse supragranular cortex. Single nucleus RNA-seq analysis of human MTG identified five glutamatergic t-types (referred to in shorthand by their most selective gene marker) with somata predominantly located in L2 and/or L3 ^11^. Similar single cell RNA-seq analysis of mouse VISp and ALM only identified three glutamatergic L2/3 t-types in each region that are known to be intratelencephalically projecting (IT) ^22^. Quantitative cross-species alignment mapped the three mouse L2/3 t-types (*Adamts2*, *Agmat* and *Rrad*) to three of the human t-types (*LTK*, *GLP2R* and *FREM3*); as mentioned, we therefore refer to these types as homologous t-types^11^. The other two human t-types (*CARM1P1* and *COL22A1*) were found in deep L3. Though all five human t-types mapped to the intratelencephalically projecting (IT) mouse subclass, consistent with the finding that supragranular cortex is composed solely of corticocortical- and telencephalon-projecting neurons, surprisingly these deep L3 human types were more similar transcriptomically to infragranular L5 and L6 IT types in mouse ^11^.

Here we extend this result to directly compare transcriptomic heterogeneity of supragranular glutamatergic neurons between mouse and human. This can be visualized using Uniform Manifold Approximation and Projection (UMAP) for dimension reduction ^28^, where the distance between cells approximates overall differences in gene expression, and consequently cells from the same t-type group together (Fig. 1d-e). In human, the overall distribution forms an extended continuum across the *LTK*, *GLP2R*, *FREM3* and *CARM1P1* t-types, with *COL22A1* cells located on a separate island (Fig. 1d), while similar analysis of the mouse L2/3 types showed much more compact distribution (Fig. 1e). As reported previously ^11^, the largest t-type, *FREM3*, showed a particularly extended graded distribution that could be split into depth-dependent subtypes with more lenient clustering (Fig. 1d; top right panel) and that varied as a function of cortical layer (Fig. 1d; bottom right panel). This within-type heterogeneity can be quantified by comparing variance explained by the first principal component (PC) in real versus shuffled data, while accounting for the number of cells in each type. This analysis confirms high heterogeneity in *FREM3*, with lower, generally similar, values for all other human and mouse t-types (Fig. 1f). To complement this analysis, we calculated the distinctness (or discreteness) between clusters as the number of differentially expressed (DE) genes between pairs of types. Homologous types are similarly discrete from one another in both mouse in human, whereas the deep L3 *CARM1P1* and *COL22A1* t-types had many more DE genes when compared to the homologous t-types (Fig. 1g). Together these results show similar levels of gene expression variability between glutamatergic neurons in human L2 and superficial L3 and mouse L2/3, with additional within-type variation and distinctive cell types in deep L3 in human.

### Patch-seq pipeline for human neurosurgical tissue analysis

To measure the electrophysiological and morphological properties of living human neurons, it is essential to use vital tissue from neurosurgical resections. A number of human studies ^12,14,15,29,30^ have found that surgically excised human neocortical tissues can be extracted, sliced, and maintained long enough to perform slice patch clamp experiments (all recordings typically take place within ~12 hours of resection, and in some cases much later ^31^). Critically, prior work established that human MTG t-types are consistently identified in both post-mortem and neurosurgically resected tissue, making this a suitable platform to establish the correspondence between morpho-electric and transcriptomic cell types. Thus, we developed a robust technology platform to apply the Patch-seq method ^23,24,25^ to acute slice preparations from human neurosurgically resected cortical tissues (Fig. 2a), and targeted pyramidal neurons from L2 and L3. Patch-seq allowed us to record from individual neurons while simultaneously filling each neuron with biocytin for subsequent imaging and morphological reconstruction. At the end of each experiment, the nucleus of the neuron was captured and processed for RNA-seq, resulting in a collective readout of single-cell electrophysiology, morphology and transcriptome modalities (Fig. 2b).

**Figure 2:**
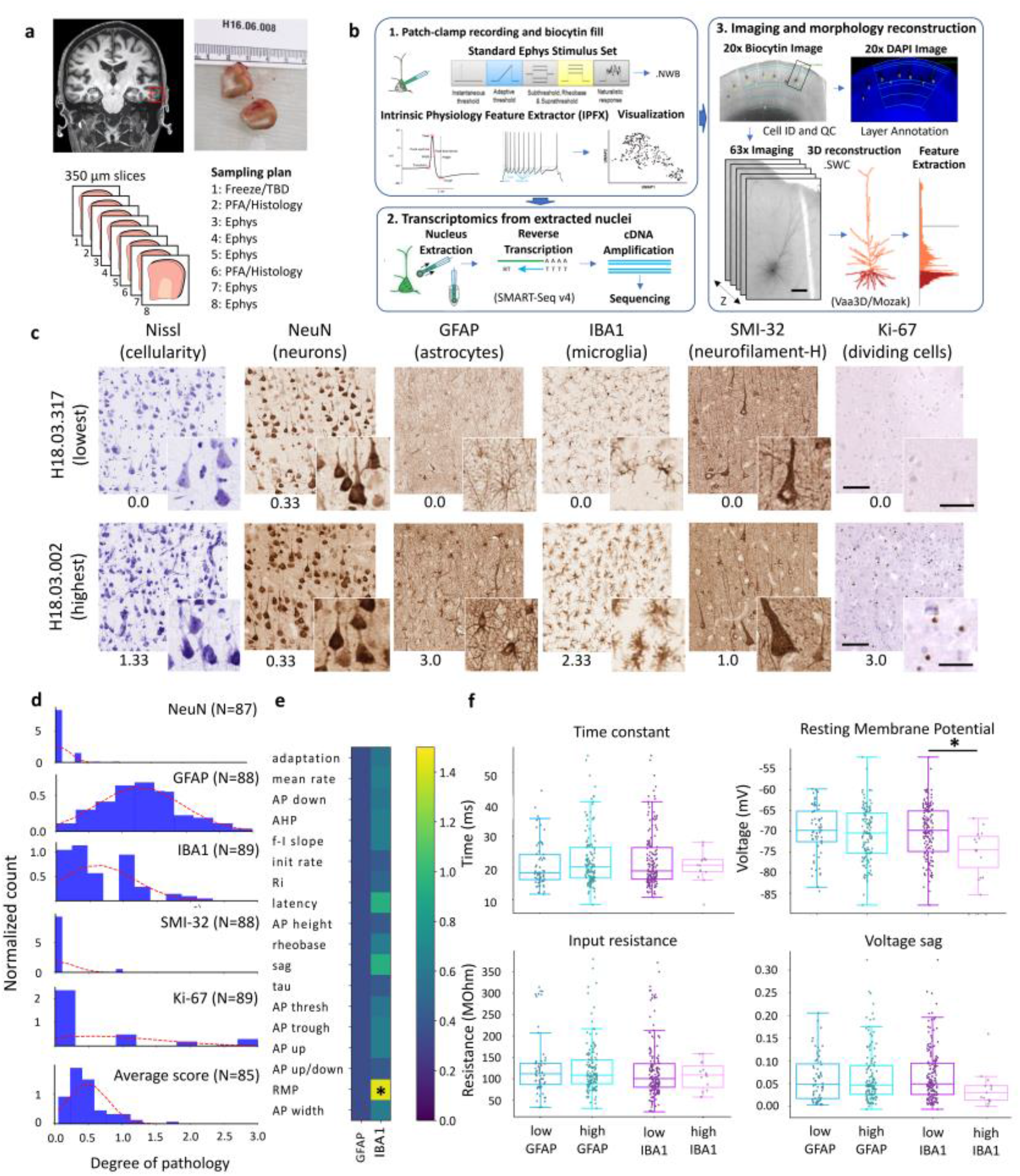
Human Patch-seq pipeline workflow and quantitative histological assessment of surgical tissue specimens. **a**) Example resected tissue specimen from human middle temporal gyrus is processed into a series of 350 μm-thick slices according to a standardized sampling plan. **b**) Workflow for patch clamp recording using standardized stimulus protocols and feature extraction code (1), followed by RNA-seq on extracted nucleated patches (2). Slices are stained with DAPI and biocytin-filled neurons are visualized with DAB as chromogen, imaged, and digitally reconstructed for morphological feature calculation and analysis (3). **c**) Immunohistochemistry and imaging on human surgical specimens using a panel of cellular markers, as indicated. Images were scored independently by three neuropathologists (from 0-3, where 0 is normal and 3 is most pathological), and the scores were averaged. Full image series are shown for the donor with the lowest (top) and highest (bottom) average marker score, demonstrating the range of cases in the study. Insets reveal cellular details. Scale bars: 50 μm for all insets and 100 μm for larger panels. Individual marker scores are indicated below each image. **d**) Histograms for four cellular markers across all donors, and for the average score across all markers in aggregate by donor (N=number of cases). **e**) Summary of statistical analysis comparing calculated electrophysiological features from recorded neurons in low (0-1) vs. high (1-3) score bins for GFAP and IBA1 cellular markers. P-values are shown as −log10(p-value). **f**) Summary plots of four selected electrophysiological features: Time constant (tau), Resting Membrane Potential (RMP), Input Resistance (Ri), and Voltage sag (sag) comparing neurons in low vs. high score bins for GFAP (blue/cyan) and IBA1 (purple/pink). Asterisk (*) indicates p<0.05 (Bonferoni corrected). Boxes show median and quartiles, whiskers show trimmed range without outliers >1.5 IQR beyond quartiles. Individual neuron data points horizontally jittered for clarity. GFAP low: 60 neurons; GFAP high: 125 neurons; IBA1 low: 170 neurons and IBA1 high: 15 neurons.

Neurons were filtered based on a series of quality-control (QC) steps for each modality (see Methods). A total of 385 neurons that passed transcriptomic data QC mapped with higher confidence to the five supragranular human glutamatergic t-types than to any other neuron type. Most neurons in the dataset preserved enough labeling to determine the relative depth of the soma with respect to the pia or the L1 - L2 border. Most neurons (283) also had sufficiently complete recordings to calculate electrophysiological features. The subset of neurons (109) with sufficient biocytin labeling and intact apical dendrites were imaged at high resolution, then subsequently manually reconstructed based on the image. L2/3 pyramidal neurons from mouse visual cortex were analyzed using the same Patch-seq platform ^32^, resulting in 120 neurons with high-quality electrophysiology and transcriptome data mapping to the three L2/3 glutamatergic t-types, and 60 neurons with data in all three modalities.

### Detailed histological assessment of human surgical tissue

A major impediment in the field regarding utilization of human neurosurgical tissue for functional studies has been the implicit assumption that patient-derived tissue specimens are inherently pathological or unhealthy, even for resected tissue distantly located from the pathological focus, thereby precluding basic discoveries about the healthy human brain. Yet prior studies have provided evidence to challenge this assumption, suggesting that surgically-resected cortical tissue slices were healthy and that pyramidal neuron morphology and physiology were largely comparable for tumor-derived versus epilepsy-derived tissue specimens, indicating the absence of overt disease-specific cellular pathology ^12,14,31^. To address this important issue further, we established a platform for comprehensive histological assessment of human surgical cases and performed an independent quantitative analysis to test for correlations or alterations in neuronal properties with respect to cellular markers of pathology.

We included several well-established histological markers for evaluating cellularity (Nissl), neuronal density and layer orientation (NeuN), astrogliosis (glial fibrillary acidic protein, GFAP), microglial activation state (IBA1), non-phosphorylated neurofilament-H (using antibody SMI-32), and cellular proliferation (Ki-67). Immunostained tissue sections were digitized, compiled by donor, and independently scored by three neuropathologists using a 4-point scale, where 0 is normal and 3 is overtly pathological (see Methods). To show the range of surgical tissue cases included in this study, images from the case with the lowest average score (mean of 6 marker scores = 0.06) are contrasted with images from the case with the highest average score (mean of 6 marker scores = 1.83) (Fig. 2c). In the highest-scored case, cellular abnormalities were clear, including astrogliosis, microglial activation, and the presence of Ki-67+ cells. However, this was rare, and most cases had average scores <1.0 (Fig. 2d), a range considered not overtly pathological. In addition, we found very low correlation of the 6 cellular markers with each other, with only Ki-67 and Nissl (cellularity) being modestly correlated (Extended Data Fig. 1). Taken together, the low correlation among the various markers and low range of average scores indicate a lack of pathology for the vast majority of surgically-resected cortical tissue samples in this study.

To assess the relationship between pathological scores and physiological properties, we binned all supragranular glutamatergic neurons derived from cases with average scores between 0-1 (low) and 1-3 (high) and directly compared the electrophysiological features of neurons with low versus high scores. For every neuron we calculated 18 electrophysiological features including input resistance (Ri), membrane time constant (tau), spike frequency adaptation (adaptation), voltage sag, resting membrane potential (RMP), and various spike-related features (Fig. 2e). Comparisons were made only if there were at least 10 neurons in low and high groups for each cellular marker. As such, our analysis was limited to GFAP and IBA1 markers (the markers with the widest spread of scores). Other markers such as SMI-32 and NeuN were highly skewed toward 0, such that all neurons derived from these markers would fall into the low bin, precluding further analysis. Among the 18 electrophysiological features analyzed for GFAP and IBA1, only one feature (resting membrane potential, RMP) related to IBA1 was significantly different between low versus high groups. Neurons in the high IBA1 group were approximately 5 mV more hyperpolarized at rest than neurons in the low IBA1 group (Fig. 2f). The remaining 17/18 features for IBA1 and all 18 features for GFAP were not statistically different between low and high groups, indicating that high scores for these specific cellular markers are overall not associated with aberrant intrinsic electrophysiological properties. This lack of association between pathology and electrophysiology can also be seen at the aggregate level in a UMAP projection of all electrophysiological features (Extended Data Fig. 2) - cells split by pathology (tumor/epilepsy) are distributed in an unstructured manner across the dataset (as are splits by other available patient characteristics including age and gender).

### High-confidence Patch-seq neuron t-type mapping

A key component of our Patch-seq approach is reliable mapping of Patch-seq cells to t-types. This issue is particularly important when analyzing individual neurons from many human individuals (potential donor-to-donor variability) undergoing neurosurgical procedures (potential disease or injury signatures). Our prior report describing the t-type classification used here ^11^ demonstrated that t-types were robust across individuals and between acute neurosurgical and postmortem frozen tissues and could be validated in independent donors with multiplex fluorescence in situ hybridization (mFISH) panels derived from these data to confirm their laminar localization. However, neurons collected via Patch-seq can exhibit contamination not seen in dissociated cells ^33^, arising from adjacent neurons and/or non-neuronal cells that enter the patch pipette. Furthermore, capturing only a portion of a neuron’s content could lead to increased variability and false negatives (dropouts). Indeed, we found much more reliable mapping when the cell nucleus was extracted, presumably due to a more consistent amount of cellular RNA and perhaps also to occlusion of the pipette opening by the nucleus against contamination.

To quantify the effect of contamination and gene dropout, we compared median gene expression levels of homologous t-types between platforms and between species (Fig. 3a). Expression data from dissociated mouse whole cells and human nuclei were moderately correlated (R=0.57, p~0, Fig. 3a, left), but with higher gene detection in whole cells, as previously shown ^21^. By comparison, dissociated nuclei and Patch-seq cells from matched human t-types were highly correlated (R=0.85, p~0, Fig. 3a, right). Relatively few genes (177 genome-wide) showed enriched expression in dissociated nuclei relative to Patch-seq cells, suggesting that high quality transcriptomes collected in this data set do not show the increased dropout rate reported in our previous study ^32^. This is likely because we are comparing our human Patch-seq cells to a reference of dissociated human nuclei, rather than dissociated mouse cells. In contrast, we identified 2,670 genes with at least four-fold enrichment in Patch-seq, including genes associated with extra-nuclear compartments such as the mitochondria (p<10^−12^) and ribosome (p<10^−9^), genes regulating cell death (p<10^−18^), RNA-binding genes (p<10^−8^) including immediate early genes, and markers for non-neuronal cells such as microglia (p<10^−20^). Some of the top genes in these categories include *COX3*, *FOS*, and *IL1B*, which all show >100-fold enrichment in Patch-seq cells. These results indicated that Patch-seq cells likely contain some RNA collected from extra-nuclear compartments and from nearby contaminating cells (particularly microglia) and may show some activity-dependent transcription. However, these effects are minor compared to species differences and we find overall high consistency and similar quality between Patch-seq cells and dissociated nuclei.

**Figure 3:**
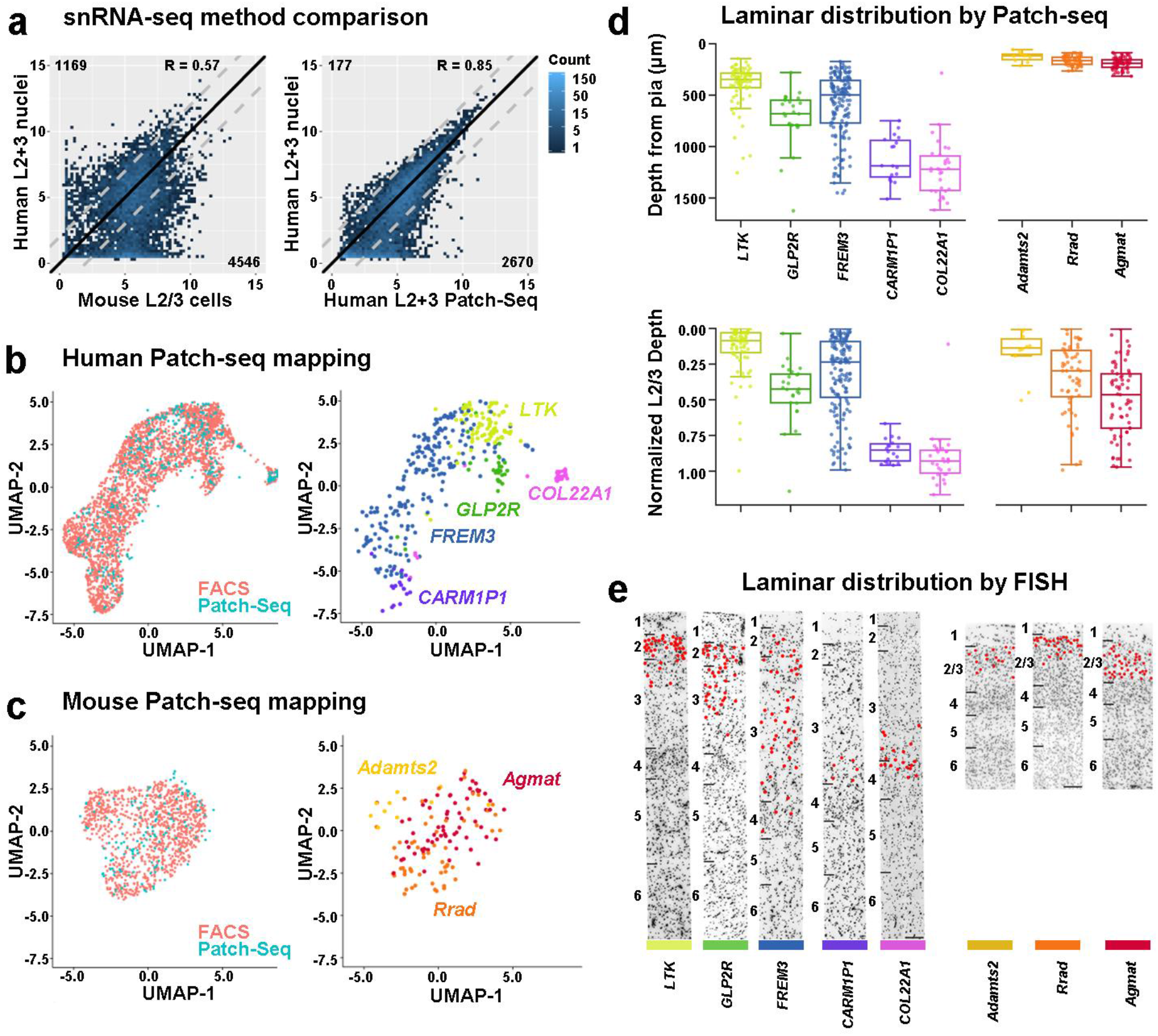
Classification of human Patch-seq neurons from supragranular cortex based on transcriptomics. **a**) Density scatter plot showing the average expression of homologous genes between the three human and mouse homologous types in dissociated cells and nuclei (left), and between dissociated nuclei and Patch-seq cells in human (right). Dashed lines indicate two-fold enrichment, with number of DE genes shown in the off-diagonal corners. p~0 for both plots. **b**) Joint UMAP of dissociated human nuclei from **Fig. 1d** and 385 glutamatergic Patch-seq neurons from supragranular cortex in MTG. Patch-seq neurons are classified using Seurat and then displayed in the same UMAP space as dissociated nuclei. Left plot shows cells color-coded by collection strategy. Right plot shows only Patch-seq neurons color-coded by mapped t-type. **c**) Joint UMAP of dissociated mouse cells from **Fig. 1e** and 133 glutamatergic Patch-seq neurons from supragranular cortex in VISp, which were classified as described for mouse GABAergic neurons ^32^. Panels and labels as in **b**. **d**) Depth distribution of neurons in human and mouse supragranular cortex, grouped and colored by t-type. Top plot shows depth from pia in μm. Bottom plot shows scaled depth within L2/3. Boxes show median and quartiles, whiskers show trimmed range without outliers >1.5 IQR beyond quartiles. Individual neuron data points horizontally jittered for clarity. **e**) Location of t-types within the cortex (indicated by red dot) demonstrated using multiplex FISH. Layer boundaries indicated by black lines. Mouse cortex is aligned to human cortex at the L1/L2 border. T-type is indicated below each image along with t-type specific color bar. Scale bar: 100 μm.

Data alignment methods have been developed to match t-types across conditions where variability across data sets is dramatically higher than variability between t-types ^34,35,36^. These strategies have been successful for comparisons of cells in different cortical regions or even in different species using the t-type classifications used in the current study ^11,37,22^. Here, human Patch-seq cells were mapped using the cell type classification workflow in Seurat (V3) ^34,35^, after first filtering out genes potentially associated with the undesirable sources of variation described above (also see Methods). Additionally, many neurons patched in mouse (but not human) supragranular cortex co-expressed GABAergic and glutamatergic genes; therefore, mapping of mouse neurons included an additional filtering step requiring expression of intronic reads that map to glutamatergic t-types as well as use of an extended reference data set (Methods). After alignment, Patch-seq cells intermix with dissociated cells and nuclei in a low-dimensional UMAP projection space in both human (Fig. 3b) and mouse (Fig. 3c), and cells assigned to the different t-types are generally co-localized in distinct locations in this space, indicating good agreement between platforms.

Biocytin staining facilitated identification of the precise cortical depth for each neuron and demonstrated a clear sublaminar distribution for each t-type in human L2 and L3 and in mouse L2/3 (Fig. 3d). In human, these sub-laminar Patch-seq distributions were remarkably consistent with histological mFISH-based spatial t-type distributions (Fig. *3*e; described previously for L3 and L4 t-types ^11^) and layer dissections in the original studies that used dissociated cells and nuclei ^11,22^. Depth distributions were also generally consistent between Patch-seq and mFISH for the three L2/3 glutamatergic t-types in mouse (Fig. *3*e). Human *LTK* neurons were found primarily in L2 and in the border region of L2 and L3. *GLP2R* neurons were found primarily in upper L3, with some neurons found in L2. *FREM3* neurons spanned L2 and L3, continuing into L4, consistent with their heterogeneous gene expression profile. *CARM1P1* and *COL22A1* were found almost entirely in deep L3 and along the L3/L4 border. Mouse L2/3 transcriptomic types also overlapped within sub-laminar space. *Rrad* and *Adamts2* neurons were located closer to the L1-L2/3 border, while *Agmat* neurons were more broadly distributed and had a greater frequency in deep L2/3. Collectively, these results indicate that Patch-seq data are consistent with reference classifications from dissociated cells or nuclei and that mapping can be robust despite many potential sources of technical noise and technical variation in human cases.

### Increased morpho-electric specialization in human L2-3 t-types

We first analyzed the three homologous L2 and L3 human and L2/3 mouse t-types, focusing on two main questions. Are there distinguishing morpho-electric phenotypes of t-types, and are they more distinct from one another in human versus mouse? The morpho-electric properties of these three t-types in aggregate were very consistent with previous reports of slice physiology recordings from human L2 and L3 pyramidal neurons ^12,13^, indicating that the Patch-seq method facilitates comparable analyses. However, with the transcriptome as the basis, human t-types showed clear qualitative morpho-electric differences (Fig. 4a; Extended Data Fig. 3). One of the most obvious differences between human t-types was cell size (e.g., dendrite height and total length), necessarily varying dramatically given the large thickness of human supragranular cortex and the laminar selectivity of different t-types with apical dendrites that extend to L1. *LTK* neurons were found primarily in L2 and upper L3. Morphologically, they were relatively short, but extended multiple apical branches into L1. Electrophysiologically, they exhibited a regular firing pattern with little firing rate adaptation and no sag. *GLP2R* neurons were found just deeper than *LTK* neurons, primarily in upper L3. They tended to have fewer dendritic branches for their longer apical extent, and often had a distinct apical tuft in L1.

**Figure 4:**
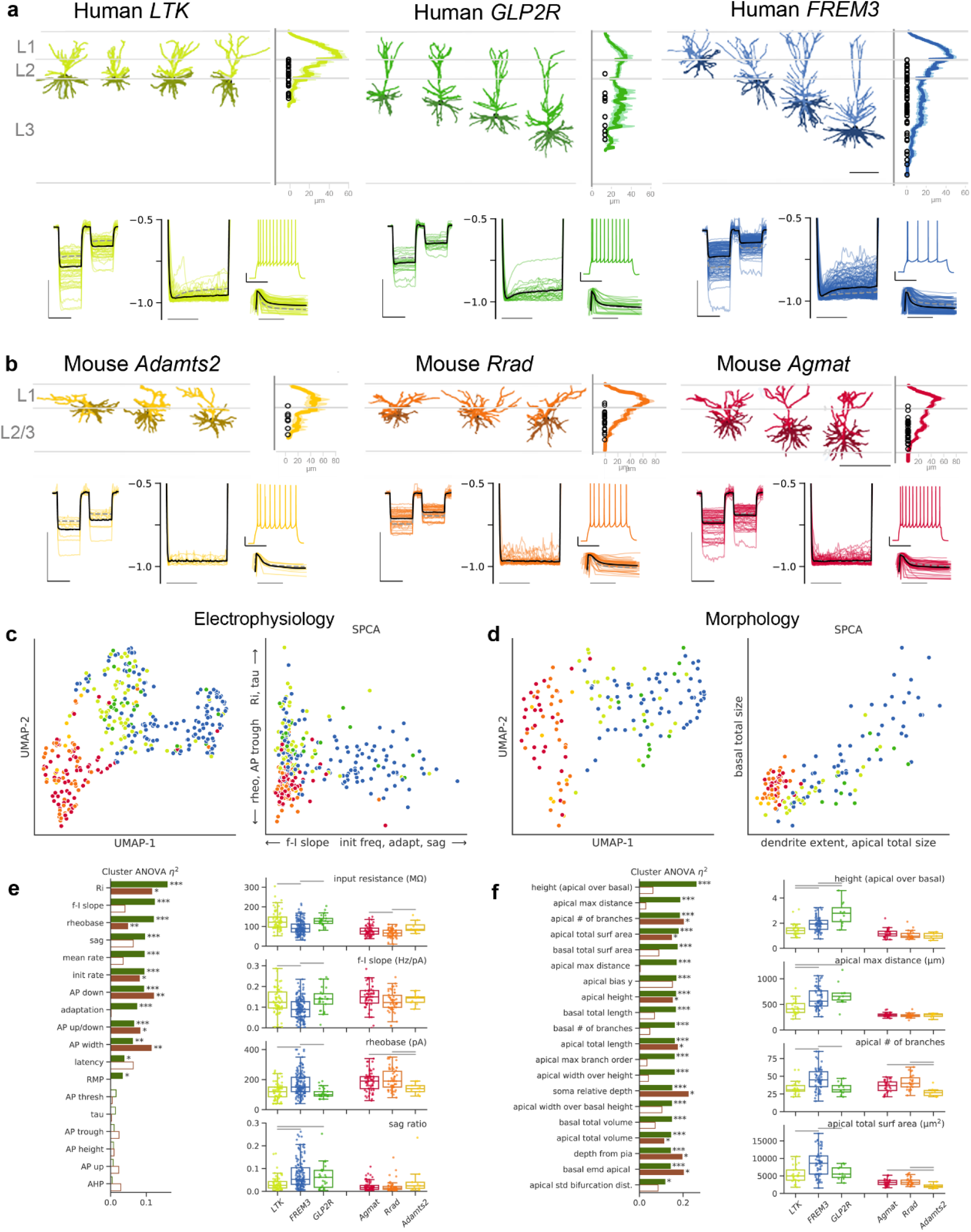
Human L2 and L3 glutamatergic t-types show greater morphological and electrophysiological differentiation than their homologous mouse L2/3 glutamatergic t-types. **a**) Morphology and electrophysiology descriptions of the three prominent human L2 and upper L3 glutamatergic neuron types: LTK, GLP2R, and FREM3. In each panel, morphology is described on top: Left, 4 representative examples of morphological reconstructions from each t-type. Scale bar = 250 μm. Right, Histogram of the average apical dendrite branch density (normalized to the maximum value for each t-type) for all reconstructed cells from each t-type. Bottom panels compare the intrinsic electrophysiological responses for 66 LTK, 25 GLP2R and 136 FREM3 neurons. For each panel, colored lines are individual neurons, solid black line represents the mean of all neurons in that t-type, dashed gray line represents the global mean of the other 2 homologous t-types in that species. Left is an overlaid response to −70 and −30 pA current injections (scale bar = 10 mV, 1.0 s), center left are hyperpolarizing pulses normalized to their peak deflection to allow for a sag comparison, shown is the range −0.5 to −1.0 (scale bar = 0.5 s). Right is a representative suprathreshold spiking response (top, scale bar = 20 mV, 0.5 s), and the normalized instantaneous firing rates for a suprathreshold pulse, demonstrating the neuron’s firing rate adaptation (bottom, scale bar = 0.5 s). **b**) Morphology and electrophysiology descriptions of the three L2/3 glutamatergic t-types in mouse visual cortex: Adamts2, Rrad, and Agmat. Panel descriptions are the same as in (A). Scale bar = 250 μm. Electrophysiological responses are shown for 9 Adamst2, 43 Rrad and 55 Agmat cells. **c**) and **d**) UMAP representation of electrophysiology and morphology space (left in each panel) generated from calculated features in each modality. Right panel in each shows the same feature space projected onto sparse principal components (SPCA), with contributing features listed on each axis. **e**) and **f**) Effect size (explained variance) for one-way ANOVA of each electrophysiology (**e**) and morphology (**f**) feature vs. t-type for human (green) and mouse (red). Stars indicate significance at FDR (False Discovery Rate) < (0.05, 0.01, 0.001). Boxplots on right show data distribution by t-type for the four features with the largest effect size in human. Gray bars indicate significant pairwise comparisons (p<0.05, FDR-corrected Mann-Whitney test). Boxes show median and quartiles, whiskers show trimmed range without outliers >1.5 IQR beyond quartiles. Individual neuron data points horizontally jittered for clarity.

Electrophysiologically, *GLP2R* neurons exhibited *LTK*-like electrophysiology, lacking adaptation and higher input resistance, but differed from *LTK* neurons in that they had pronounced sag. The *FREM3* t-type represented 56.7% of supragranular glutamatergic neurons collected in L2 or L3 dissections ^11^, with a laminar distribution that spanned the entire distribution of *LTK* and *GLP2R* and had morpho-electric properties that were overlapping but distinct from those t-types. *FREM3* neurons varied from small neurons in upper L2 to very large neurons in the deeper part of L3 and had a gradient of morpho-electric properties like the graded transcriptional properties described above. Upper L2 *FREM3* neurons had an apical dendrite restricted to L1 and L2 and regular firing while the large, deep L3 *FREM3* neurons had an apical dendrite that spanned L1-3 and a heavily adapting firing rate (described further below). Apical dendrites of *FREM3* and *LTK* neurons branched much closer to the soma than *GLP2R* neurons at the same depth, resulting in more radial branching across layers.

Mouse L2/3 t-types also had heterogeneous electrophysiological and morphological properties, but in general were more like one another than the human t-types (Fig. 4b; Extended Data Fig. 4). As mentioned, each of the three t-types had distributions spanning all of L2/3, although there was a trend for *Adamts2* and *Rrad* to be more superficial (Fig. *3*e). Similarly, each of these t-types contained neurons with wide and tufted branching ^2,38,39^. Since we are comparing non-homologous brain regions between species, we verified that the electrophysiological differences between L2/3 glutamatergic neurons in mouse visual cortex compared to mouse temporal association area (TEa), a region of rodent cortex previously used for human temporal cortex comparisons ^12,40,41,42^, were far smaller than those seen between mouse VISp and human temporal cortex (Extended Data Fig. 5).

**Figure 5:**
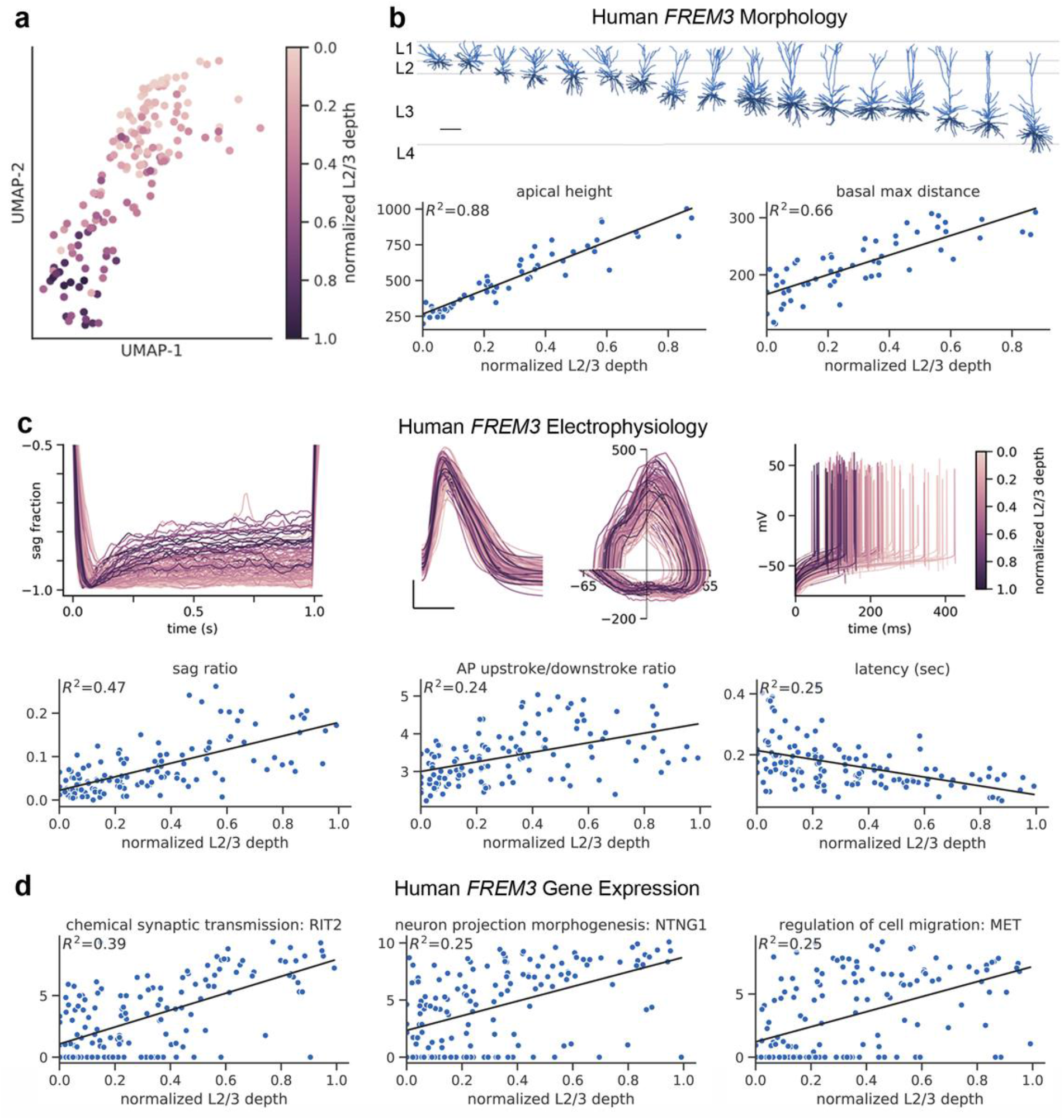
Morpho-electric-transcriptomic features of FREM3 neurons vary according to laminar depth. **a**) FREM3 neurons plotted in transcriptomic UMAP space (as in Fig. 3c). Each cell is colored based on its relative position within L2-3. Depth color scale shown at right. **b**) FREM3 neurons exhibit a range of sizes for morphologies spanning L2-3. Scale bar = 250 μm. Apical height and basal maximum (max) branch distance are positively correlated with depth. For (b-d), all regressions shown are significant at FDR<10^−7^. **c**) Top, electrophysiology data traces colored based on each neuron’s relative position within L2-3 (scale at right). Top left, hyperpolarizing pulses normalized to their peak deflection to allow for a sag comparison (N=124). Top middle, overlaid first action potential during a rheobase current injection (scale bar = 25 mV, 1.0 ms, traces aligned to the time of threshold), as well as the corresponding phase plots (x axis, mV; y axis, mV/ms). Top right, Initial action potentials at rheobase for FREM3 neurons (N = 141) aligned to the time of stimulus onset. Bottom, summary plots show in FREM3 neurons that sag and action potential (AP) upstroke / downstroke ratio are positively correlated with depth while latency to AP firing at rheobase is negatively correlated with depth. **d**) Representative gene examples for three GO categories with pronounced depth dependence of expression in FREM3 neurons, chemical synaptic transmission, neuron projection morphogenesis, and regulation of cell migration.

To quantify the magnitude of differences between and within transcriptomic types, we calculated 18 electrophysiological features that characterize passive, single action potential, and suprathreshold properties as well as 60 morphological features that capture the extent and complexity of basal and apical dendrites, their distribution across cortical layers and soma position. In UMAP representations of the combined human and mouse electrophysiology and morphology data (Fig. 4 c and d), human and mouse neurons occupy separate islands, reflective of the number and magnitude of differences in morpho-electric properties (Extended Data Table 1). A sparse principal component analysis (SPCA) projection of the electrophysiological features (Fig. 4c right) was used to select small groups of features that determine two axes of greatest variability across the dataset. The first (y-axis) is dominated by features largely related to passive properties (e.g., dendrite surface area and membrane composition), including membrane time constant and input resistance. Variability along the second (x-axis) principally differentiates between human neurons and is explained by properties like adaptation rate and sag that are less clearly related to neuron size. Likewise, an SPCA projection of morphological features (Fig. 4d right) shows that features related to the total size of apical dendrites (length, volume, and surface area) and the spatial extent of apical and basal dendrites best explain the species differences, while additional basal dendrite size features capture further variability between human neuron t-types.

To quantify the degree of separation or overlap in different features between mouse and human t-types, we ran a one-way ANOVA for the effect of t-type on each calculated electrophysiology and morphological feature. For electrophysiological features (Fig. 4e), 3/18 showed differences between t-types that explained >10% of feature variance (*R*^2^>0.1) for both human and mouse (FDR<0.05 for mouse features, <10^−5^ for human). However, the mouse types were distinct in input resistance and two related AP shape features (width and downstroke) with a maximum R^2^ =0.12, while the human types showed distinct firing properties (f-I slope and rheobase) in addition to input resistance, with a maximum R^2^ =0.16. For morphological features (Fig. 4f), 16/60 features had R^2^ >0.15 between the human t-types (FDR<10^−3^), compared to 12/60 for the mouse t-types (with 10/12 significant at FDR<0.05). This quantitative analysis confirms the qualitative observation that the main supragranular human t-types are more morpho-electrically specialized from one another than their mouse homologues, primarily in terms of morphology, but with a moderate contrast in electrophysiology as well.

Finally, we also quantified to what degree this morpho-electric differentiation can be used to differentiate among the human t-types and mouse t-types, training a random forest classifier to predict t-type identity using electrophysiological or morphological features. Homologous human t-types were predicted with 67% accuracy using electrophysiological features, while mouse t-types were predicted with 54% accuracy. Classifying based on morphological and electrophysiological features combined resulted in slightly improved performance of 69% for human neurons and 60% for mouse neurons (Extended Data Fig. 6). This moderate cross-species contrast in predictability reinforces the contrast in t-type differentiation from ANOVA, although the overall accuracy is low, limited in part by low cell numbers for some human and mouse t-types. Additionally, in both analyses the between-type variability in the human t-types may be partially obscured by the significant variability within the *FREM3* t-type, as discussed below.

**Figure 6 (previous page):**
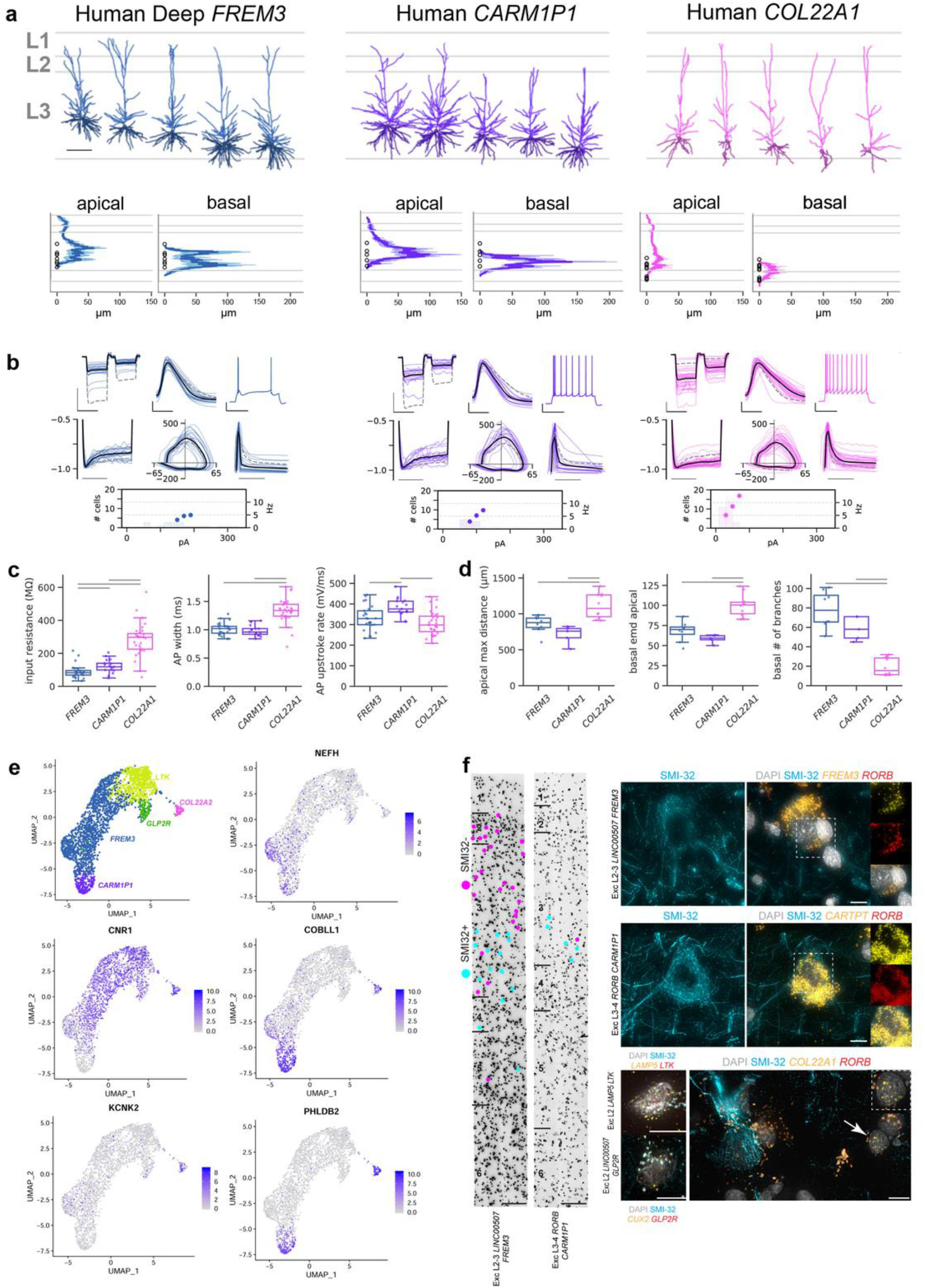
Human deep L3 glutamatergic t-types are morphologically and electrophysiologically distinct. **a**) Top panels, representative morphological reconstructions of the two deep human L3 glutamatergic t-types, CARM1P1 and COL22A1 neurons, compared to the deep FREM3 neurons. Scale bar = 250 μm. Bottom panels, histogram of average apical dendrite branch length (normalized to the maximum value for each t-type) for all reconstructed neurons from each t-type. **b**) Electrophysiological description of the intrinsic electrophysiology responses of 21 deep FREM3, 17 CARM1P1 and 37 COL22A1 neurons. For each panel, colored lines are individual cells, solid black line represents the mean of all neurons in that t-type, dashed gray line represents a global mean across the other two deep human t-types. Top left is an overlaid response to −70 and −30 pA current injections (scale bar = 10 mV, 1.0 s); bottom left are hyperpolarizing pulses normalized to their peak deflection to allow for a sag comparison (scale bar = 0.5 s), shown is the range −0.5 to −1.0. Center top shows overlaid first action potential during a rheobase current injection (scale bar = 25 mV, 1.0 ms), center bottom is the corresponding phase plot (x axis, mV; y axis, mV/ms). Top right is a representative suprathreshold spiking response (scale bar = 20 mV, 0.5 s), and bottom right are the normalized instantaneous firing rates for a suprathreshold pulse, demonstrating adaptation of firing rate (scale bar = 0.5 s). Bottom: histogram of rheobase current injections (left axis) and frequency to current relationships for each neuron (right axis), normalized to the mean rheobase current for each t-type. **c** and **d**) Summary of electrophysiology (**c**) and morphology (**d**) features that discriminate the three deep t-types from each other. Features shown were selected from significant ANOVA results (FDR<10^−7^ in c, FDR<10^−2^ in d). Gray bars indicate significant pairwise comparisons (p<0.05, FDR-corrected Mann-Whitney test). Boxes show median and quartiles, whiskers show trimmed range without outliers >1.5 IQR beyond quartiles. Individual neuron data points horizontally jittered for clarity. **e**) A selection of five marker genes that are differentially expressed in the deep L3 human t-types. **f**) SMI-32 immunostained MTG tissue. FREM3 and CARM1P1neurons that are SMI-32ir indicated by pink dots and those not SMI-32ir indicated by blue dots. Layer boundaries indicated at left of image, Scale bar = 100 μm. Representative SMI-32 immunoreactivity photomicrographs, along with mFISH for t-type specific genes shown for FREM3 (top) and CARM1P1 (middle) t-types. At right, representative mFISH composite images showing labeling for DAPI, neurofilament H, CARTPT and RORB in the same cell. White box indicates region of image shown at right where CARTPT and RORB are shown separately and then combined. At bottom, representative mFISH composite images showing labeling for DAPI, neurofilament H, and t-type-specific genes for LTK t-type (LAMP5 and LTK), GLP2R t-type (CUX2 and GLP2R) and COL22A1 t-type (COL22A1 and RORB) Scale bar = 10 μm. Marker gene expression in Extended Data Fig. 10).

### Human FREM3 t-type exhibits depth-dependent morpho-electro-transcriptomic variation

As described above, the *FREM3* t-type displays graded features as a function of cortical depth. Anatomically, *FREM3* neurons span the full depth of L2 and L3 and send apical dendrites to L1, at distances of > 1 mm, which is greater than the entire thickness of mouse VISp (Fig. *3*d, 5b). Transcriptomically, *FREM3* is also the most heterogeneous t-type among the human L2/3 glutamatergic t-types, exhibiting graded gene expression that correlates with the cell’s inferred laminar location based on relatively coarse laminar dissections (Fig. 1d and ^11^). Patch-seq laminar positions confirm this depth vs. transcriptome relationship directly. UMAP plots of the *FREM3* neuron population generated based on transcriptomic data reveal a clear relationship between soma depth and gene expression (Fig. 5a). Superficial neurons with somata in L2 and upper L3 (<500 μm from the border of L2 and L3) mostly appear at the top of the transcriptomic UMAP space and transition gradually into neurons with somata located 500-1000 μm from this border (Fig. 5a). Deep L3 neurons (1500 μm) all appear at the bottom of the transcriptomic UMAP space with a partial separation between the superficial and deep sub-regions. Similarly, multiple electrophysiological and morphological features (apical and basal dendrites) vary continuously with depth (Fig. 5b, c) in agreement with previous findings ^13,14,12^. Graded change in morphological phenotype across the cortical depth is shown in the representative morphological reconstructions in Figure 5b. Graded change in electrophysiological properties, including sag and action potential latency, is shown in the raw traces colored by soma depth in Figure 5c.

To explore the full range of depth-related variation across all data modalities, we calculated correlations with normalized L2/3 depth for each electrophysiological, morphological and gene feature. Nine out of 18 physiological features were significantly correlated with depth (FDR<0.05, 7/18 FDR<0.01; Supplementary Data 1). The three strongest electrophysiology correlations with depth were the increase in sag and AP (Action Potential) upstroke / downstroke ratio and the decrease in AP latency at rheobase (Fig. 5c). 37 out of 58 morphological features were correlated with depth (FDR<0.05, 28/58 FDR<0.01). While features like apical height are effectively constrained to follow the distance from the soma to L1, many functionally independent features were also strongly correlated, including the maximum length of basal dendrites (Fig. 5b) and soma radius (Extended Fig. 11; Supplementary Data 1). 790 genes were correlated with depth (FDR<0.05) and GO (gene ontology) enrichment analysis on this gene set revealed a variety of significantly enriched functional categories (Fig. 5d, Supplementary Data 1) that predict functional variation in different neuronal phenotypes. Graded genes were enriched for genes associated with synaptic transmission, and developmental processes like cell migration and neuron projection morphogenesis. For example, the receptor tyrosine kinase gene *MET* is implicated in neuronal growth, synaptic function and cortical circuit function, while Netrin-G1 (*NTNG1*) is involved in axonal and dendritic outgrowth associated with specific circuit formation. These molecular differences suggest differences in neuronal connectivity in *FREM3* neurons as a function of their laminar position.

### Deep L3 human glutamatergic t-types are morpho-electrically distinct

The *CARM1P1* and *COL22A1* human t-types do not have homologous types in mouse supragranular cortex, although they were shown to be most like glutamatergic IT types in deeper infragranular layers of mouse cortex ^11^. This could be interpreted as a species difference in cellular migration of conserved types to different laminar positions; however, the deeper L3 *FREM3* neurons taken alone also map best to infragranular mouse IT t-types (data not shown). Rather, there appears to be an overall shift in transcriptomic similarity by depth of human neurons to mouse neurons such that deeper L3 is more like L5 and L6 in mouse, and *CARM1P1* and *COL22A1* t-types likely represent evolutionarily new types in human (and at least other primate species (Bakken et al., bioRxiv 2020)). This finding is consistent with previous work ^43^ that uncovered a set of genes showing a dramatic shift from L5 expression in mouse cortex to expression in large L3 pyramidal neurons in human temporal and visual cortex.

The morpho-electric properties of *CARM1P1* and *COL22A1* neurons differed markedly from the human L2 and L3 homologous types (Fig. 4; Extended Data Fig. 7) and each other (Fig. 6 a-d). Though *CARM1P1* and *COL22A1* neurons co-mingle with the largest *FREM3* neurons and also have large somata (Extended Data Fig. 8), they are restricted to the deepest part of L3 where they form a highly diverse set of putative IT projection neuron types. In order to understand how *CARM1P1* and *COL22A1* differ from these deep *FREM3* neurons specifically, we split the *FREM3* t-type by depth, with the neuronal density minimum in L3 as a dividing line (L2/3 depth = 0.575; Fig. 1c), and directly compared the morpho-electric properties of these deep types. In contrast to deep *FREM3* neurons, the *CARM1P1* t-type exhibited extensive proximal apical oblique and basal dendritic branching (Fig. 6a). In fact, *CARM1P1* neurons had the largest total dendritic length of all the L2 and L3 t-types, despite having a shorter apical dendrite length on average. Electrophysiologically, *CARM1P1* neurons exhibited a faster action potential upstroke than the other deep types (Fig. 6c). This extensive dendritic branching did not predict a lower input resistance, which was higher than deep *FREM3* neurons (Fig. 6c).

*COL22A1* differed notably from *CARM1P1* and deep *FREM3* neurons. *COL22A1* had very sparse dendritic branching (Fig. 6a). The primary apical dendrite branched near the soma and extended just one or two branches into superficial layers. Minimal L1 branching was a consistent feature of the deepest L3 glutamatergic neurons across multiple t-types. *COL22A1* neurons exhibited very high input resistance (Fig. 6c) and thus were the most responsive to current injection, displaying a steeper firing frequency to current input gain relative to the other deep L3 t-types. Interestingly, *COL22A1* neurons showed a smaller amount of sag than *CARM1P1* neurons (Fig. 6b) located at an equivalent distance from pia, indicating that this property is t-type-specific rather than depth-specific.

Repeating the ANOVA analysis of electrophysiological and morphological features with these three deep types showed that as a group they are quantitatively as well as qualitatively more distinct than the three human homologous L2 and L3 types described in Fig. 4. ANOVA differences were seen in 15 out of 18 electrophysiological features and 24 out of 60 morphological features (FDR<0.05) (Supplementary Data 1). Many of these features showed very large effect sizes, with the variance explained by type surpassing 40% (*R*^2^>0.4) for 4 electrophysiological and 20 morphological features, while no features met this threshold for the three homologous types (maximum *R*^2^ =0.26).

To understand transcriptional differences that may be predictive of phenotypic differences between *CARM1P1*, *COL22A1*, and deep *FREM3* neurons, we used genesorteR ^44^ with slightly relaxed parameters (quant = 0.7) to identify DE genes selective for one or two of these three t-type sets, and found 219 such marker genes (Extended Data Fig. 9). Since dissociated nuclei were not collected using sublaminar dissections, deep *FREM3* neurons were defined as *FREM3* neurons dissected from L3 or L4 that were assigned to subtype f73 (Fig. 1), which colocalizes with deep *FREM3* Patch-seq neurons in UMAP space (Fig. 3c, 5a). Furthermore, 77 of these 219 marker genes (including four genes shown in Fig. 6e) were also defined as marker genes by Patch-seq, where cortical depth was explicitly measured, suggesting the selection of deep *FREM3* neurons in dissociated nuclei was reasonable.

Differences in morpho-electric properties of the three deep L3 t-types were reflected in DE genes enriched for GO terms associated with neuronal connectivity, structure, and synaptic signaling, including axon (p=3.5 10^−6^; Bonferroni corrected), synapse (p=5.3 10^−5^), calcium ion binding (p=0.008), and extracellular matrix organization (p=0.00002). For example, cannabinoid receptor type 1 (*CNR1*) is highly expressed in the *COL22A1* t-type but not in the *CARM1P1* t-type (Fig. 6e), implying a cell-type specific difference in the effects of cannabinoid compounds. In contrast, both *PHLDB2* and *COBLL1* are highly expressed in *COL22A1* and *CARM1P1* t-types, with *PHLDB2* displaying slightly more specific enrichment of expression in these t-types. *COBLL1* is a morphogenesis-associated gene that has been shown to promote dendrite branching and the formation of actin filament membrane ruffles ^45^. Similarly, *PHLDB2* localizes to dendritic spines of hippocampal neurons where it plays an important role in regulation of long-term potentiation by affecting the density of glutamate receptors, and knockout of this gene impairs the formation of memories in mice ^46^. *KCNK2*, which shows relatively selective expression in *COL22A1* neurons, is a potassium channel, which can convert between voltage-insensitive potassium leak current and voltage-dependent outward rectifying current depending on phosphorylation ^47^, and knockdown of this gene in mice impairs the migration of late-born neurons destined to become glutamatergic neurons in L2/3 ^48^.

Differential connectivity patterns of glutamatergic neurons have been described in L3 of macaque temporal cortex based on immunolabeling for the SMI-32 antibody that recognizes non-phosphorylated epitopes of the neurofilament heavy chain ^16^. SMI-32ir neurons preferentially make long-range ipsilateral projections, whereas neurons that are not SMI-32ir tend to make more proximal local projections ^16^. Furthermore, the SMI-32ir L3 neurons show selective vulnerability in AD ^17,18^. These observations in monkey are consistent with our finding that connectivity-related genes vary between t-types. To assess the relationship between deep L3 t-type and projection phenotypes we combined SMI-32 immunoreactivity with mFISH for markers of *FREM3*, *CARM1P1* and *COL22A1* neurons (Fig. 6f). The large *FREM3* and *CARM1P1* neurons were SMI-32ir, while *COL22A1* neurons were not SMI-32ir. Furthermore, the gene coding SMI-32, *NEFH,* also shows increased expression in deep *FREM3* and *CARM1P1* relative to *COL22A1* and all the superficial glutamatergic t-types (Fig. 6e). This finding creates a putative link between transcriptomically-defined cell types, long-range target specificity, and vulnerable neuron populations in AD.

## Discussion

Understanding the fundamental cellular components of cortical circuits has been a major goal of neuroscience from the time of Ramón y Cajal ^49^. However, a robust, quantitative, widely agreed upon definition of cell types and delineation of cellular diversity has been elusive due to the high degree of cellular complexity and low-throughput techniques available for cellular analysis that lead to underpowered statistical analyses. This challenge is compounded in human cortex, where limited access to tissue, high variation across individuals, and lack of reliable genetic tools have severely limited progress. Recent advances in single-cell or single-nucleus transcriptomics have revolutionized our understanding of cellular diversity. Recent studies involving tens of thousands of cells from single cortical regions in mouse and human have derived cellular classifications, based on similar patterns of gene expression, that appear to mirror many aspects of cellular cytoarchitecture, function, and developmental origins ^22,11^. In principle, the transcriptome represents the complete set of genes coding for cellular phenotypes, but for the most part the high degree of neuronal diversity described by transcriptomics analyses remains to be validated as meaningful by demonstrating correlation with other structural and functional properties. Furthermore, transcriptomics provides evidence both for discrete cell classes as well as more continuous variation within cell classes ^11,22,50,51^ whose functional relevance has not yet been demonstrated in the adult nervous system. This transcriptomic landscape provides a powerful framework for bounding the problem of cellular diversity, allowing targeted analysis of transcriptomically-defined cell types coupled with analysis of other neuronal phenotypes using techniques such as the triple modality Patch-seq method used here.

A consistent critique of cellular and molecular studies using neurosurgically resected tissues is that there must be huge variation associated with disease state and neuropathology that will obscure any coherent results. Indeed, many studies have used human surgical tissue from the pathological focus to identify disease-related phenomena ^52,53,54,55,56^. However, a growing number of studies have shown highly consistent results using neocortical tissues distal to the sites of frank pathology ^14,57,15,29,12,13^. To rigorously explore this variability, in the current study we implemented a standardized histological analysis with markers for neurons, astrocytes and microglia to look for neuron loss, glial proliferation or inflammatory responses, along with blinded neuropathologist scoring. Importantly, we found little evidence for consistent disease- or pathology-related alterations of the physiological features measured when using cortical tissues from MTG (predominantly epilepsy cases) or frontal cortex regions (predominantly tumor cases) with no obvious gross pathology. These findings indicate that typical cellular properties can be robustly studied in surgically resected human neocortical tissues. Indeed, a remarkable result from the current study is the stereotypy of neuron types, measured transcriptomically, morphologically and physiologically, across human neurosurgical specimens from 90 different tissue donors with many uncontrolled axes of variation such as age, gender, ethnicity, disease condition and severity. This indicates that the basic cellular blueprint is highly robust across individuals and can be studied routinely using surgically-excised tissues from hospitals around the world. The magnitude of differences observed between human and mouse here highlight the importance of taking advantage of such clinical tissues to gain a strong understanding of the details of human brain functional organization in health and disease.

A principal result of the current study is that the transcriptomes of neurons in human supragranular cortex are well-correlated with morphological and physiological features, as well as cortical depth. Many morpho-electric features vary between transcriptomic cell types. The *LTK* t-type contains the smallest neurons and is largely restricted to L2, while the *GLP2R* t-type is found in L2 and superficial L3 with fewer dendrites and minimal branching in L2. The *FREM3* t-type found throughout L2 and the full depth of L3 has small to large neurons across this depth. The *CARM1P1* and *COL22A1* t-types are highly distinctive and found exclusively in deep L3 with wide, highly branched apical dendrites and tall, very sparse apical dendrites, respectively. In addition, the most abundant *FREM3* t-type shows strong continuous variation as a function of cortical depth transcriptomically; this molecular variation is correlated with continuous variation of the morphological and physiological features of these neurons. For example, apical and basal dendritic length both increase with depth. Multiple other physiological features also vary with depth such as input resistance and sag. Together these results suggest that the transcriptome serves as something of a Rosetta stone for understanding supragranular glutamatergic neurons and reveals several organizational principles. First, morpho-electro-transcriptomic neuron types occupy different depths in supragranular cortex, which has more diversity than previously described. Second, cortical layers are enriched for specific neuron types, but are also highly heterogeneous with multiple neuron types that cross laminar boundaries. Finally, continuous variation of a single (*FREM3*) t-type through the full ~1mm-depth of human supragranular cortex is a major axis of functional organization.

New analytical techniques aligning transcriptomic datasets ^34,58^ have enabled mouse and human transcriptomic databases to be related ^11^. These analyses indicated a general conservation of cortical cell types but with substantial species differences. In supragranular cortex, neurons from all three mouse and the three most abundant human glutamatergic types all mapped to a single cross-species superset, rather than at the finest level of resolution in either species. Neuronal diversity increased in deep L3 and L4 (whose boundaries are not sharp), and these human types mapped either to mouse L4 types, or, surprisingly, to deep layer mouse neurons. Importantly, in mouse the relationship between axonal projection class and transcriptomic types has been established, and the distinction between IT neuron types and locally or deep subcortically projecting neurons is very robust. All the human deep L3 types map to IT types, suggesting that (as in mouse and monkey) the neurons that make up supragranular cortex are all part of the IT class.

Comparative analysis of the anatomical and physiological properties of mouse and human supragranular neurons substantiate and extend the previous transcriptomic results and illustrate that the evolutionary expansion of supragranular cortex is accompanied by many changes in glutamatergic neuron types. These differences can be summarized as 1) increased phenotypic differentiation of conserved transcriptomic types, 2) increased degree of graded variation as a function of depth in the cortex within the most abundant type, and 3) increased neuronal diversity with addition of new types in deep L3.

On the first point, a prominent result predicted by cytoarchitecture is that the main IT types that make up L2 and L3 in human are much more different from one another than their L2/3 homologues in mouse. One aspect of this is simply the anatomical positioning of cell bodies, which has become more spread out in the ~1 mm depth of human L2 and L3 compared to the ~250 μm depth of mouse L2/3. There is an easily definable L2 in human cortex where the majority of L2 *LTK* neurons are located, whereas *GLP2R* neurons extend deeper into L3, and *FREM3* neurons are found throughout L2 and L3. As discussed above, many other physiological features and anatomical features vary among these human neuron types; in contrast, although there is some variation in depth within L2/3, mouse supragranular neurons appear to be quite homogeneous and very few features differ among the three mouse types.

The second notable difference between mouse and human is the high degree of depth-dependence for features of the most abundant *FREM3* t-type compared to mouse. A variety of anatomical features vary as a function of depth in this type, and the variance as a whole can be quite large; for example, the maximum basal dendritic path length varies from 100 to 300 μm from L2 to the deepest part of L3, and the soma diameter from 8 to 28 μm. Physiological features, including sag, latency, and spike shape (upstroke/downstroke ratio), also vary as a function of depth and differ significantly between mouse and human. These features may work in concert with previously reported distinctive human intrinsic membrane properties to mediate between-species differences in spatial-temporal synaptic integration ^15,40,42,12^.

Finally, the deep part of human L3 contains a greater diversity of neuron types than observed in mouse L2/3. As described previously using only transcriptomics, these neurons map best to mouse L5 and L6 infragranular IT neurons ^11^. As described here, these two t-types, *CARM1P1* and *COL22A1*, are highly distinctive transcriptomically, anatomically, and physiologically. The *CARM1P1* neurons are very large with profuse basal and oblique dendrites, similar to the largest *FREM3* neurons in deep L3 but most often with apical dendrites that conspicuously do not reach L1. The *COL22A1* neurons are very different, with elongated somas and simple untufted apical dendrites that frequently do not reach L1. How should this species difference be interpreted? At least two plausible explanations exist. One possibility is that these human supragranular types are homologous to the best transcriptomically matching mouse infragranular types but have migrated to different cortical locations in development. This idea is supported by transcriptomic similarity and earlier observations that many mouse infragranular layer genes are instead found predominantly in human supragranular layers ^43^. In this interpretation, this increased supragranular neuron diversity represents an anatomical (and functional) reorganization of the cortical microcircuit. Another possibility is that these represent evolutionarily distinct neuron types that are similar to mouse infragranular IT types. Because this homology alignment is based on transcriptomic similarity, new types that have coopted preexisting transcriptional programs will appear similar. Indeed, even the deeper half of the conserved *FREM3* t-type, if mapped to mouse types independently from the more superficial *FREM3* neurons, align best with deep layer mouse IT neuron types. A parsimonious explanation may come from developmental biology. The cortex is generated in an inside-out fashion with neurons destined for more superficial layers generated sequentially from a common progenitor pool over time. Transcriptomically, this developmental sequence is reflected in the adult, with neurons in adjacent layers showing greatest similarity; single cell transcriptomics has extended this to show a general depth-dependence to similarities that may reflect developmental origin. Human excitatory neuron corticogenesis is dramatically extended compared to mouse, and it could be that the overall progression along this developmental trajectory is shifted such that the expanded L2 and L3 occur at different times along this trajectory.

By whichever mechanism, the outcome is that the deeper part of L3 contains a greater diversity of IT neurons in human relative to mouse cortex. Neurons in this area have long been an area of focused study in non-human primate species, where large deep L3 pyramidal neurons have been shown to selectively express (along with pyramidal neurons in L5 and L6) the non-phosphorylated form of heavy chain neurofilament protein ^16,59^. Neurons in the non-human primate expressing this protein in L3c are known to make long-range, predominantly ipsilateral projections compared to more locally projecting neurons. We show here that the deeper L3 *FREM3* and *CARM1P1* neurons, but not the *COL22A1* neurons, express the mRNA for *NEFH* and label with antibody SMI-32 in human L3. SMI-32 immunolabeling has been shown to be depleted in L3 magnopyramidal neurons in AD progression ^18,17^, indicating a selective vulnerability of the largest long-range association neurons and consequent disruption of cortical networks affected in AD pathology. The current results add to this finding by showing that neurofilament-H maps onto the transcriptomic, morphological, and physiological classification, labeling some types but not others. This refined cellular perspective serves as a new roadmap for future studies investigating selective neuron vulnerability and resistance, and for exploring the functional implications of loss of those connections in AD.

## Methods

Detailed descriptions of all experimental data collection methods in the form of technical white papers can also be found under ‘Documentation’ at http://celltypes.brain-map.org.

### Human tissue acquisition

Surgical specimens were obtained from local hospitals (Harborview Medical Center, Swedish Medical Center and University of Washington Medical Center) in collaboration with local neurosurgeons. All patients (Extended Data Table 2) provided informed consent and experimental procedures were approved by hospital institute review boards before commencing the study. Tissue was placed in slicing artificial cerebral spinal fluid (ACSF) as soon as possible following resection. Slicing ACSF was comprised of (in mM): 92 N-methyl-D-glucamine chloride (NMDG-Cl), 2.5 KCl, 1.2 NaH_2_PO_4_, 30 NaHCO_3_, 20 4-(2-hydroxyethyl)-1-piperazineethanesulfonic acid (HEPES), 25 D-glucose, 2 thiourea, 5 Na-L-ascorbate, 3 Na-pyruvate, 0.5 CaCl_2_.4H_2_O and 10 MgSO_4_.7H_2_O ^60^. Prior to use, the solution was equilibrated with 95% O_2_, 5% CO_2_ and the pH was adjusted to 7.3 by addition of 5N HCl solution. Osmolality was verified to be between 295-305 mOsm/kg. Human surgical tissue specimens were immediately transported (15-35 min) from the hospital site to the laboratory for futher processing.

### Mouse breeding and husbandry

All procedures were carried out in accordance with the Institutional Animal Care and Use Committee at the Allen Institute for Brain Science. Animals (<5 mice per cage) were provided food and water ad libitum and were maintained on a regular 12 hour light–dark cycle. Animals were maintained on the C57BL/6J background, and newly received or generated transgenic lines were backcrossed to C57BL/6J. Experimental animals were heterozygous for the recombinase transgenes and the reporter transgenes.

### Tissue processing

For mouse experiments, male and females were used between the ages of P45 and P70 were anesthetized with 5% isoflurane and intracardially perfused with 25 or 50 ml of 0-4°C slicing ACSF. Human or mouse acute brain slices (350 μm) were prepared with a Compresstome VF-300 (Precisionary Instruments) or VT1200S (Leica Biosystems) vibrating microtome modified for block-face image acquisition (Mako G125B PoE camera with custom integrated software) before each section to aid in registration to the common reference atlas. Brains or tissue blocks were mounted for slicing with the optimal orientation for preserving intactness of apical dendrites of cortical pyramidal neurons.

Slices were transferred to an oxygenated and warmed (34 °C) slicing ACSF for 10 min, then transferred to room temperature holding ACSF of the composition (in mM): 92 NaCl, 2.5 KCl, 1.2 NaH_2_PO_4_, 30 NaHCO_3_, 20 HEPES, 25 D-glucose, 2 thiourea, 5 Na-L-ascorbate, 3 Na-pyruvate, 2 CaCl_2_.4H_2_O and 2 MgSO_4_.7H_2_O ^60^ for the remainder of the day until transferred for patch-clamp recordings. Prior to use, the solution was equilibrated with 95% O_2_, 5% CO_2_ and the pH was adjusted to 7.3 using NaOH. Osmolality was verified to be between 295-305 mOsm/kg.

### Patch-clamp recording

Slices were bathed in warm (32-34°C) recording ACSF containing the following (in mM): 126 NaCl, 2.5 KCl, 1.25 NaH_2_PO_4_, 26 NaHCO_3_, 12.5 D-glucose, 2 CaCl_2_.4H_2_O and 2 MgSO_4_.7H_2_O (pH 7.3), continuously bubbled with 95% O_2_ and 5% CO_2_. The bath solution contained blockers of fast glutamatergic (1 mM kynurenic acid) and GABAergic synaptic transmission (0.1 mM picrotoxin). Thick-walled borosilicate glass (Warner Instruments, G150F-3) electrodes were manufactured (Narishige PC-10) with a resistance of 4–5 MΩ. Before recording, the electrodes were filled with ~1.0-1.5 μL of internal solution with biocytin (110 mM potassium gluconate, 10.0 mM HEPES, 0.2 mM ethylene glycol-bis (2-aminoethylether)-N,N,N′,N′-tetraacetic acid, 4 mM potassium chloride, 0.3 mM guanosine 5′-triphosphate sodium salt hydrate, 10 mM phosphocreatine disodium salt hydrate, 1 mM adenosine 5′-triphosphate magnesium salt, 20 μg/ml glycogen, 0.5U/μl RNAse inhibitor (Takara, 2313A) and 0.5% biocytin (Sigma B4261), pH 7.3). The pipette was mounted on a Multiclamp 700B amplifier headstage (Molecular Devices) fixed to a micromanipulator (PatchStar, Scientifica).

The composition of bath and internal solution as well as preparation methods were made to maximize the tissue quality, to align with solution compositions typically used in the field (to maximize the chance of comparison to previous studies), and modified to reduce RNAse activity and ensure maximal recovery of mRNA content.

Electrophysiology signals were recorded using an ITC-18 Data Acquisition Interface (HEKA). Commands were generated, signals processed, and amplifier metadata were acquired using MIES (*https://github.com/AllenInstitute/MIES/*), written in Igor Pro (Wavemetrics). Data were filtered (Bessel) at 10 kHz and digitized at 50 kHz. Data were reported uncorrected for the measured (Neher 1992) –14 mV liquid junction potential between the electrode and bath solutions.

Prior to data collection, all surfaces, equipment and materials were thoroughly cleaned in the following manner: a wipe down with DNA away (Thermo Scientific), RNAse Zap (Sigma-Aldrich) and finally with nuclease-free water.

Neuron targeting: For human slices, pyramidal shaped neurons in L2 and 3 were targeted to recording. For mouse experiments, pyramidal neurons in L2/3 were targeted, either tdTomato-pyramidal neurons when recording from a transgenic line that labels interneurons, or tdTomato+ neurons when recording from a line that labels different populations of L2/3 glutamatergic neurons, specifically Oxtr-T2A-Cre and Penk-IRES2-Cre-neo, each crossed to the Ai14 tsTomato reporter line.

After formation of a stable seal and break-in, the resting membrane potential of the neuron was recorded (typically within the first minute). A bias current was injected, either manually or automatically using algorithms within the MIES data acquisition package, for the remainder of the experiment to maintain that initial resting membrane potential. Bias currents remained stable for a minimum of 1 s before each stimulus current injection.

To be included in analysis, a cell needed to have a >1GΩ seal recorded before break-in and an initial access resistance <20 MΩ and <15% of the Rinput. To stay below this access resistance cut-off, cells with a low input resistance were successfully targeted with larger electrodes. For an individual sweep to be included, the following criteria were applied: (1) the bridge balance was <20 MΩ and <15% of the Rinput; (2) bias (leak) current 0 ± 100 pA; and (3) root mean square noise measurements in a short window (1.5 ms, to gauge high frequency noise) and longer window (500 ms, to measure patch instability) <0.07 mV and 0.5 mV, respectively.

Upon completion of electrophysiological examination, the pipette was centered on the soma or placed near the nucleus (if visible). A small amount of negative pressure was applied (~-30 mbar) to begin cytosol extraction and attract the nucleus to the tip of pipette. After approximately one minute, the soma had visibly shrunk and/or the nucleus was near the tip of the pipette. While maintaining the negative pressure, the pipette was slowly retracted in the x and z direction. Slow, continuous movement was maintained while monitoring pipette seal. Once the pipette seal reached >1GΩ and the nucleus was visible on the tip of the pipette, the speed was increased to remove the pipette from the slice. The pipette containing internal solution, cytosol and nucleus was removed from pipette holder and contents were expelled into a PCR tube containing the lysis buffer (Takara, 634894).

### cDNA amplification and library construction

We performed all steps of RNA-processing and sequencing as described for mouse Patch-seq cells ^32^. We used the SMART-Seq v4 Ultra Low Input RNA Kit for Sequencing (Takara, 634894) to reverse transcribe poly(A) RNA and amplify full-length cDNA according to the manufacturer’s instructions. We performed reverse transcription and cDNA amplification for 20 PCR cycles in 0.65 ml tubes, in sets of 88 tubes at a time. At least 1 control 8-strip was used per amplification set, which contained 4 wells without cells and 4 wells with 10 pg control RNA. Control RNA was either Universal Human RNA (UHR) (Takara 636538) or control RNA provided in the SMART-Seq v4 kit. All samples proceeded through Nextera XT DNA Library Preparation (Illumina FC-131-1096) using either Nextera XT Index Kit V2 Sets A-D(FC-131-2001,2002,2003,2004) or custom dual-indexes provided by IDT (Integrated DNA Technologies). Nextera XT DNA Library prep was performed according to manufacturer’s instructions except that the volumes of all reagents including cDNA input were decreased to 0.2x by volume. Each sample was sequenced to approximately 1 million reads.

### RNA sequencing data processing

Fifty-base-pair paired-end reads were aligned to GRCh38.p2 using a RefSeq annotation gff file retrieved from NCBI on 11 December 2015 for human and to GRCm38 (mm10) using a RefSeq annotation gff file retrieved from NCBI on 18 January 2016 for mouse (*https://www.ncbi.nlm.nih.gov/genome/annotation_euk/all/*). Sequence alignment was performed using STAR v2.5.3 ^61^ in two pass Mode. PCR duplicates were masked and removed using STAR option “bamRemoveDuplicates”. Only uniquely aligned reads were used for gene quantification. Gene counts were computed using the R Genomic Alignments package summarizeOverlaps function using “IntersectionNotEmpty” mode for exonic and intronic regions separately ^62^. Expression levels were calculated as counts of exonic plus intronic reads. For most analyses, log2(counts per million [CPM] + 1) transformed values were used.

### Morphological Reconstruction

#### Biocytin histology

A horseradish peroxidase (HRP) enzyme reaction using diaminobenzidine (DAB) as the chromogen was used to visualize the filled cells after electrophysiological recording, and 4,6-diamidino-2-phenylindole (DAPI) stain was used identify cortical layers as described previously ^2^.

#### Biocytin labeled neuron imaging

Mounted sections were imaged as described previously ^2^. Briefly, operators captured images on an upright AxioImager Z2 microscope (Zeiss, Germany) equipped with an Axiocam 506 monochrome camera and 0.63x optivar. Two-dimensional tiled overview images were captured with a 20X objective lens (Zeiss Plan-NEOFLUAR 20X/0.5) in brightfield transmission and fluorescence channels. Tiled image stacks of individual cells were acquired at higher resolution in the transmission channel only for the purpose of automated and manual reconstruction. Light was transmitted using an oil-immersion condenser (1.4 NA). High-resolution stacks were captured with a 63X objective lens (Zeiss Plan-Apochromat 63x/1.4 Oil or Zeiss LD LCI Plan-Apochromat 63x/1.2 Imm Corr) at an interval of 0.28 μm (1.4 NA objective) or 0.44 μm (1.2 NA objective) along the Z axis. Tiled images were stitched in ZEN software and exported as single-plane TIFF files.

#### Morphological reconstruction

Reconstructions of the dendrites and the full axon were generated for a subset of neurons with good quality transcriptomics, electrophysiology and biocytin fill. Reconstructions were generated based on a 3D image stack that was run through a Vaa3D-based image processing and reconstruction pipeline ^63^. Images were used to generate an automated reconstruction of the neuron using TReMAP (Zhou 2016). Alternatively, initial reconstructions were created manually using the reconstruction software PyKNOSSOS (Ariadne-service) or the citizen neuroscience game Mozak (Mosak.science)^64^. Automated or manually-initiated reconstructions were then extensively manually corrected and curated using a range of tools (e.g., virtual finger, polyline) in the Mozak extension (Zoran Popovic, Center for Game Science, University of Washington) of Terafly tools ^65,66^ in Vaa3D. Every attempt was made to generate a completely connected neuronal structure while remaining faithful to image data. If axonal processes could not be traced back to the main structure of the neuron, they were left unconnected.

Before morphological feature analysis, reconstructed neuronal morphologies were expanded in the dimension perpendicular to the cut surface to correct for shrinkage ^13,67^ after tissue processing. The amount of shrinkage was calculated by comparing the distance of the soma to the cut surface during recording and after fixation and reconstruction. A tilt angle correction was also performed based on the estimated difference (via CCF registration) between the slicing angle and the direct pia-white matter direction at the cell’s location ^2^.

### Slice Immunohistochemistry

#### Immunohistochemistry

Tissue slices (350 μm-thick) designated for histological profiling were fixed for 2-4 days in 4% paraformaldehyde (PFA) in phosphate-buffered saline (PBS) at 4°C and transferred to PBS + 0.1% sodium azide for storage at 4°C. Slices were then cryoprotected in 30% sucrose, frozen and re-sectioned at 30 μm using a sliding microtome (Leica SM2000R). Sections were stored in PBS+azide at 4°C in preparation for immunohistochemical and Nissl staining. Specific probes (vendor, dilution) used were: Neu-N (Millipore #MAB377, 1:2000); SMI-32 (Biolegend #801704, 1:2000); GFAP (Millipore #MAB360, 1:1500); Parvalbumin (Swant #PV235, 1:2000); Iba-1 (Wako #019-19741, 1:1000); Ki67 (Dako #M724001-2, 1:200). Full immunohistology protocol details available at *http://help.brain-map.org/download/attachments/8323525/CellTypes_Morph_Overview.pdf?version=4&modificationDate=1528310097913&api=v2*

#### Slide imaging

Colorimetric IHC and other histologically-stained whole slides (i.e. Nissl-stained preparations) for brightfield imaging were scanned using an Aperio ScanScope XT slide scanner (Leica Biosystems, Germany). The samples were illuminated using a 21DC Halogen Lamp (Techniquip, USA). Brightfield images were acquired using ScanScope Console (v101.0.0.18) and controller (ve101.0.4.446) at 10x magnification (objective lens 20x/0.75 NA Plan Apo, 0.5x magnifier) resulting in a pixel size of 1.0 μm/pixel.

### Multiplex fluorescent in situ hybridization (FISH)

Fresh-frozen human postmortem brain tissues were sectioned at 14-16 μm onto Superfrost Plus glass slides (Fisher Scientific). Sections were dried for 20 minutes at −20°C and then vacuum sealed and stored at −80°C until use. The RNAscope multiplex fluorescent v1 kit was used per the manufacturer’s instructions for fresh-frozen tissue sections (ACD Bio), except that fixation was performed for 60 minutes in 4% paraformaldehyde in 1X PBS at 4°C and protease treatment was shortened to 10 minutes. For combined RNAscope and immunohistochemistry, primary antibodies were applied to tissues after completion of mFISH staining. Primary mouse anti-Neurofilament H (SMI-32, Biolegend, 801701) was applied to tissue sections at a dilution of 1:250. Secondary antibodies were goat anti-mouse IgG (H+L) Alexa Fluor conjugates (594 or 647). Sections were imaged using a 60X oil immersion lens on a Nikon TiE fluorescence microscope equipped with NIS-Elements Advanced Research imaging software (version 4.20). For all RNAscope mFISH experiments, positive cells were called by manually counting RNA spots for each gene. Cells were called positive for a gene if they contained ≥ 3 RNA spots for that gene. Lipofuscin autofluorescence was distinguished from RNA spot signal based on the larger size of lipofuscin granules and broad fluorescence spectrum of lipofuscin. The following probe combinations were applied to label cell types of interest: (1 - LTK) LTK (NM_002344.5), LAMP5 (NM_012261.3); (2 – GLP2R) GLP2R (NM_004246.2), CUX2 (NM_015267.3); (3 – FREM3) RORB (NM_006914.3), FREM3 (NM_001168235.2); (4 – CARM1P1) RORB, CARTPT NM_004291.3); (5 – COL22A1) RORB, COL22A1 (NM_152888.3); (6 – Adamts2) Cbr3 (NM_173047.3), Neurod1 (NM_010894.2), Cdh13 (NM_019707.4); (7 – Rrad) Nr4a3 (NM_015743.3), Cux1 (NM_009986.4), Cdh13; (8 – Agmat) Pou3f2 (NM_008899.2), Igfbp7 (NM_001159518.1), Coch (NM_001198835). Experiments were repeated on at least N=2 donors per probe combination for both mouse and human.

### Quantification of human and mouse soma size

Images of NeuN+ stained sections from human MTG (1 section per donor for 5 donors) and mouse VISp (1 section per animal for 3 animals) (described above) were imported into ImageJ for processing. ROIs were drawn around cell bodies and exported as “.roi” files for downstream processing. In both species, L4 is defined as a band of densely packed, small granular cells, and the upper bound of this band (which includes overlying large pyramidal cells) is treated as the border between L3 and 4. The border between L1 and 2 is defined as the sharp boundary between the cell-sparse zone of L1 and the is a cell-dense zone of L2. In mouse, the border between L2 and 3 is indistinguishable and not defined. In human MTG, the boundary between L2 and 3 can be closely approximated as transition from densely packed small pyramidal cells to less densely packed larger pyramidal cells, which is largely consistent among donors.

Soma areas were defined as the number of pixels contained in each ROI, scaled by the number of pixels per μm. Cortical depth was defined for each cell as the position of that cell centroid relative to pia (absolute depth) or relative to the L1/2 and L3/4 boundaries (scaled depth) at that position in the tissue. The number of neurons per mm^2^ of L2/3 cortex (absolute density) is the number of neurons per image scaled by the area of the image where cell counts were assessed. For measuring surface density and cell area across L2/3 cortical depth, L2/3 was split into 20 evenly size bins and the relevant measurements within each bin were calculated independently per section (one section per donor) and the average and standard deviation across sections were reported. The first and last bins are omitted from plots as they display boundary effects. Relative (scaled) neuron density scales to 1 for each donor and is defined as the fraction of total neuron count in each bin. In human, a nadir of scaled density was identified at −0.575, which we define as a quantitative boundary between superficial and deep L3 in this manuscript.

### Analysis of data from dissociated cells/nuclei

Reference data used in this study include dissociated excitatory cells (mouse) or nuclei (human) collected from human MTG ^11^ and mouse VISp ^22^, and are all publicly accessible at the Allen Brain Map data portal (*https://portal.brain-map.org/atlases-and-data/rnaseq*). In human, cells from the five previously identified L2/3 glutamatergic types were retained, subsampling to match the laminar distribution of neurons included in the Patch-seq data set as closely as possible, leaving a total of 2,948 neurons from *LTK, GLP2R, FREM3, CARM1P1,* and *COL22A*1 t-types. In mouse, all neurons from the three L2/3 glutamatergic t-types (*Adamts2*, *Rrad*, and *Agmat*) were retained. Data sets were visualized as follows. First, the top 2,000 most binary genes by beta score ^11^, which is defined as the squared differences in proportions of cells/nuclei in each cluster that expressed a gene above 1, normalized by the sum of absolute differences plus a small constant (∊) to avoid division by zero. Scores ranged from 0 to 1, and a perfectly binary marker had a score equal to 1. Second, the Seurat pipeline ^34,35^ (more details below) was used to scale the data, reduce the dimensionality using principal component analysis (30 PCs). These PCs were then used to generate a Uniform Manifold Approximation and Projection (UMAP) ^28^. Finally, data and metadata such as cluster, subcluster, layer, and gene expression are then overlaid onto this UMAP space with using different colored or shaded points.

Cluster heterogeneity is defined as average observed variance explained by the first PC compared with permuted data after accounting for differences in the number of cells per cell type. To get this, we i) randomly selected 80 cells from each cell type, ii) identified the 80 most variable genes using the “FindVariableFeatures” Seurat function with selection.method=”vst”, iii) performed PCA after removing outlier cells, iv) calculated the percent of variance explained by the first PC, v) repeated i-iv for 100 sets of data where the expression levels for each gene are shuffled across the 80 cells to break gene correlations but retain other gene statistics, and vi) identify the average and standard deviation of PC1 for observed vs permuted data. Cluster discreteness is defined as the average number of DE genes between a given type and each of the remaining homologous t-types (*LTK*, *GLP2R*, and *FREM3* t-types in human; *Rrad*, *Agmat*, and *Adamts2* t-types in mouse). In this case pairwise differential expression is defined using the de_score function in the “scrattch.hicat” R library ^22^ after subsampling each cluster to 80 cells, and only the genes with higher expression in the relevant cluster are considered. The getMarkers function from the genesorteR R library (*https://github.com/mahmoudibrahim/genesorteR*) ^44^ was used to identify genes differentially expressed genes between deep FREM3 (f73 subtype; collected from L3 or L4 dissection), COL22A1, and CARM1P1 neurons, using all default parameters except quant=0.7. To validate the cell selection for deep L3 (since sublaminar dissection was not performed on the dissociated nuclei data), this analysis was repeated on Patch-seq neurons from these three types collected in deep L3 (scaled depth > −0.575). Gene ontology (GO) enrichment analysis was performed using ToppGene ^68^ with default settings, and Bonferroni corrected p-values are reported unless stated otherwise.

### Dataset curation

Patch-seq cells were included in this data set if they met the following criteria. All neurons: 1) had high-quality transcriptomic data, measured as the normalized summed expression of “on”-type marker genes (NMS, adapted from the single-cell quality control measures in ^33^) greater than 0.4; and 2) retained a soma through biocytin processing and imaging such that an accurate laminar association could be made. In addition, mouse neurons were: 1) located within VISp; 2) either tdTomato- or tdTomato+ from a line known to label glutamatergic neurons (i.e. tdTomato+ neurons from known inhibitory mouse lines were excluded); 3) mapped to L2/3 IT VISp Rrad, L2/3 IT VISp Agmat, or L2/3 IT VISp Adamts2 using Seurat mapping (as described below); and 4) mapped to L2/3 IT VISp Rrad, L2/3 IT VISp Agmat, L2/3 IT VISp Adamts2, or L4 IT VISp Rspo1 in a separate Seurat mapping analysis where only reads located within gene introns are considered for both data sets. This final filter removes Patch-seq cells that jointly express markers for GABergic and glutamatergic cells, likely representing L2/3 GABAergic neurons contaminated with adjacent glutamatergic cells. We do not find examples of such cells in human, possibly due to a much smaller sampling of GABAergic cells than in the mouse.

### Identifying transcriptomic types

Due to the differences in gene expression between Patch-seq and dissociated cells (see Fig. 3a and ^32^), we used transcriptomes of dissociated human nuclei from ^11^ or cells from ^22^ as reference dataset for human and mouse, respectively, and mapped Patch-seq transcriptomes to the reference data to identify their cell types. Prior to data transfer, we filtered genes potentially related to technical variables. X- and Y-chromosomes were excluded to avoid nuclei mapping based on sex. Many mitochondrial genes have expression correlated with RNA-seq data quality in dissociated nuclei data ^11^, so nuclear and mitochondrial genes downloaded from Human MitoCarta2.0 ^69^ were excluded as well. We also find that Patch-seq cells often have high expression of non-neuronal marker genes, so any genes most highly expressed in a non-neuronal cell type are excluded. Finally, any genes showing at least four-fold higher expression in dissociated nuclei vs. Patch-seq cells in the included cell types (or vice versa) were excluded as potentially platform dependent. In total 23,129 of 50,281 genes (46%) remained in human and a comparable fraction for mouse. Variable genes for mapping were selected as described above for dissociated nuclei data visualization, by using the top 2000 remaining genes by beta score as input into the procedure described below.

For both species, we mapped Patch-seq data sets to the relevant dissociated cells or nuclei reference using Seurat V3 (*https://satijalab.org/seurat/*) ^34,35^ following the tutorial for Integration and Label Transfer with default parameters for all functions, except when they differed from those used in the tutorial, and replacing variable gene selection with the genes described above. More specifically, we first define a low (30) dimensional PCA space of the dissociated cells or nuclei data set and then project this onto the Patch-seq data set. We then found transfer anchors (cells that are mutual nearest neighbors between data sets) in this subspace. Each anchor is weighted based on the consistency of anchors in it’s local neighborhood, and these anchors were then used as input to guide label transfer (or batch-correction), as proposed previously ^70^. We then scaled the data, reduced the dimensionality using principal component analysis, and visualized the results with Uniform Manifold Approximation and Projection (UMAP) ^28^. This process is done using the “FindTransferAnchors” and “TransferData” R functions, which provides both the best mapping cell type and a confidence score. For mouse data, the three homologous types did not provide a heterogenous enough reference data set, and therefore a larger set of glutamatergic and GABAergic cell types was used as reference. Cell type assignments for most cells were robust to choice of reference data set and to changes in parameter settings. Some cells with expression levels intermediate to two cell types changed calls between different runs; however, the cell type-level results presented are robust to these small changes.

Gene expression of Patch-seq cells was visualized by projection into the UMAP space calculated from dissociated nuclei using a combination of Seurat and the R implementation of the “umap” library (*https://github.com/tkonopka/umap*). More specifically, the Seurat data integration pipeline (functions “FindIntegrationAnchors” and “IntegrateData”) was used to calculated a scaled data for both data sets and PCA was performed on this integrated space. The first 30 PCs from both data sets, as well as the UMAP coordinates calculated for dissociated nuclei above were input into the umap pipeline and the “predict” function was used to project the Patch-seq cells into UMAP coordinates. As above, data and meta-data were then overlaid on these umap coordinates.

### Comparison of gene expression between species

Gene orthologs between mouse and human were pulled from the gene orthologs table on NIH (*https://ftp.ncbi.nlm.nih.gov/gene/DATA/gene_orthologs.gz*) on 22 November 2019. Only genes with unique orthologs between mouse and human were included in cross species analyses.

### Electrophysiology feature analysis

Electrophysiological features were measured from responses elicited by short (3 ms) current pulses and long (1 s) current steps as previously described (*Gouwens et al., 2019*). Briefly, APs were detected by first identifying locations where the smoothed derivative of the membrane potential (dV/dt) exceeded 20 mV/ms, then refining based on several criteria including threshold-to-peak voltage and time differences and absolute peak height. For each AP, threshold, height, width (at half-height), fast after-hyperpolarization (AHP), and interspike trough were calculated (trough and AHP measured relative to threshold), along with maximal upstroke and downstroke rates dV/dt and the upstroke/downstroke ratio (i.e., ratio of the peak upstroke to peak downstroke). Additional features from supratheshold sweeps included the rheobase and slope of the firing rate vs. current curve (f-I slope); the first spike latency initial firing rate (inverse of first ISI), measured at rheobase; and the mean firing rate and spike frequency adaptation ratio (mean ratio of consecutive ISIs), measured at ~50 pA above rheobase. Subthreshold features included the resting membrane potential (RMP), time constant (tau) from responses to short pulses, input resistance from responses across hyperpolarizing long steps, and sag ratio from response at ~ −100 pA. All feature calculation used the IPFX package (Intrinsic Physiology Feature Extraction, *https://github.com/AllenInstitute/ipfx*).

### Morphology feature analysis

Morphological features were calculated as previously described^2^. Briefly, feature definitions were collected from prior studies^1,71^. Features were calculated using the version of neuron_morphology package (*https://github.com/alleninstitute/neuron_morphology/tree/dev*). Reconstructed neurons were aligned in the direction perpendicular to pia and white matter. Additional features, such as the laminar distribution of axon, were calculated from the aligned morphologies. Shrinkage correction was not performed (see above), features predominantly determined by differences in the z-dimension were not analyzed to minimize technical artifacts due to z-compression of the slice after processing.

### Analysis of features by t-type and species

Combined datasets of electrophysiological and morphological features across homologous t-types from mouse and human were visualized by an analysis pipeline of data imputation and standardization, followed by projection to two dimensions using UMAP or SPCA (sklearn and umap python packages) ^72,73^. Cells with more than 3/18 electrophysiological features missing were dropped, the remaining missing features were imputed as the mean of 5 nearest neighbors (“KNNImputer”), and features were centered about the median and scaled by IQR (“RobustScaler”). The SPCA regularization parameter was adjusted to minimize nonzero features while preserving dataset structure. All features with coefficients over 0.05 were reported directly in the case of electrophysiology or summarized by feature categories for morphology.

For each feature, differentiation by t-type was assessed by running a one-way ANOVA for the feature by t-type, using the statsmodels package ^74^. This analysis was repeated separately for the 3 mouse and human homologous t-types, as well as the 3 deep human t-types (with the subset of deep FREM3 cells only). Results were reported as fraction of variance explained (*η*^2^ or *R*^2^) and heteroscedasticity-robust F test p-value (“HC3”), corrected for false discovery rate (Benjamini-Hochberg procedure) across all features for each data modality. Post-hoc Mann-Whitney rank tests were run across pairs of t-types in each group (human and mouse homologous types and deep human types) for top-ranked features from ANOVA, and results FDR-corrected.

For classification of t-types, features were normalized using the standard scaler scalar in sklearn (“StandardScalar”), and the data was randomly assigned with stratification to training (70%) and testing sets (30%). The random forest classifier was trained using the sklearn package with 600 decision trees. The classification performance was estimated after averaging the results of the classifiers trained on 1000 stratified random data splits and compared against performance for data with shuffled t-type labels. Confusion matrices shown are for a single representative train/test split.

### Analysis of features by depth for FREM3 t-type

For each electrophysiology, morphology, and gene feature, the depth-related variability was assessed by a linear regression of the feature against relative L2/3 depth, using the statsmodels package ^74^. Results were reported as fraction of variance explained (*R*^2^), Pearson correlation *r*, and heteroscedasticity-robust F test p-value (“HC3”), corrected for false discovery rate (Benjamini-Hochberg procedure) across all features for each data modality. Due to the large number of morphology and genes tested, results were summarized by calculating GO term enrichment in ToppGene ^68^ for the set of depth-correlated genes (FDR<0.05), followed by subselection of representative GO terms using REViGO ^75^. Groups of features were ranked by the group’s highest *R*^2^, and the features with highest correlation shown for the top groups.

## Data and software availability

The custom electrophysiology data acquisition software (MIES) is available at *https://github.com/alleninstitute/mies*. The Vaa3D morphological reconstruction software, including the Mozak extension, is freely available at www.vaa3d.org and its code is available at *https://github.com/Vaa3D*. Code for reproducing most of the analyses presented in this work are available on GitHub *https://github.com/AllenInstitute/patchseq_human_L23*.

## Author Contributions

I.S., H.M., G.T., H.Z., C.K., E.S.L. conceptualized the project. J.T.T., N.De., A.Bel., T.Ca., P.C., R.A.D., N.H., C.H., L.Ke., C.La., D.Ma., E.Me., J.N., A.Ol., J-G.Y., P.B., C.C., P.C.d-W.H., R.G.E., M.F., R.P.G., S.I., C.D.K, A.L.K., J.G.O., A.P.P., D.L.S. contributed to neurosurgical tissue acquisition. N.De., E.B., T.Ca., P.C., K.C., M.K., G.M., A.Oz., J.S., H.T., G.T. prepared brain slices. K.Ba., S.F., A.A.G., N.G., K.H., T.S.H., D.B.H., D.H., L.Ki., R.M., E.J.M., N.M., G.M., Li.N., G.O., A.Ol., A.Oz., R.R., J.T., R.W., G.T. performed electrophysiological experiments. D.B., A.Gl., J.G., D.Mc., T.P., C.R., M.T., A.T., K.W., K.S. prepared single cell RNASeq libraries. K.Bi., J.Bo., K.Br., T.E., A.Ga., H.G., M.Max., M.Mc., L.M., V.O., D.P., C.A.P., A.R. processed slices for biocytin staining and immunohistochemistry. S.D., N.Do., R.E., M.G., M.H., K.N., P.R.N., V.O., L.P., S.R. imaged immuno-stained and biocytin-stained slices, cells. S-L.D., L.A., R.D., R.A.D., T.D., A.M.H., S.K., A.M., D.S., G.W. reconstructed neurons and/or provided anatomical annotations. J.Be., S.A.S., J.T.T., J.A.M., T.Ch., A.Buc., A.Bud., S-L.D., R.D.H., B.K., B.R.L., O.F., C.G., L.Ke., K. E. L., M.Mal., Z.Y. performed analyses. J.Be., S.A.S., J.T.T., B.K., B.R.L., P.C., E.R.T., A.Ber. contributed to methods development studies. A.Bud., T.B, L.G., T.J., D.R. generated tools for pipeline data generation. F.D., S.M., S.M.S., L.E. provided program management support. J.Be., S.A.S., N.De., B.R.L., R.D., T.D., C.F., M.Mc., P.R.N., L.P., G.J.M., H.Z. organized and managed pipeline data generation. D.F., S.G., A.S., W.W., Ly.N. organized and managed pipeline data storage and processing. J.Be., S.A.S., J.T.T., J.A.M., T.Ch., A.Buc., T.E.B., A.Bud., R.D.H., B.K., B.R.L., N.G., M-H.K., B.P.L., Z.Y., C.A., A.A., A.Ber., P.R.H., A.R.J., G.J.M., J.W.P., R.Y., I.S., C.P.J.d-K., H.M., G.T., H.Z., C.K., E.S.L. provided scientific direction. J.Be., S.A.S., J.T.T., J.A.M., T.Ch., A.Buc., A.Bud., S-L.D., R.D.H., E.S.L. prepared the figures. J.Be., S.A.S., J.T.T., J.A.M., T.Ch., A.Buc., C.A., E.S.L. wrote the manuscript in consultation with all others.

## Acknowledgements

We thank Adrian Wanner for providing reconstruction services through Ariadne and Roy Szeto and Zoran Popovic for facilating the reconstruction work contributed by Mozak.science. Finally, we thank the Mozak citizen-scientists for their valuable contribution. The research was partially supported by several grant awards from institutes under the National Institutes of Health (NIH), including award numbers R01EY023173 from The National Eye Institute and U01MH105982 from the National Institute of Mental Health and Eunice Kennedy Shriver National Institute of Child Health & Human Development to H.Z. Research reported in this publication was supported by the National Institute Of Mental Health of the National Institutes of Health under Award Number U01MH114812. The content is solely the responsibility of the authors and does not necessarily represent the official views of the National Institutes of Health. The project described was supported by award number R011EY023173 from The National Institute of Allergy and Infectious Disease. Its contents are solely the responsibility of the authors and do not necessarily represent the official views of the National Institutes of Health and The National Institute of Allergy and Infectious Disease. Work was all supported by the Hungarian Academy of Sciences, the National Research, Development and Innovation Office of Hungary GINOP-2.3.2-15-2016-00018, the Ministry of Human Capacities of Hungary 20391-3/2018/FEKUSTRAT to G.T. Neuropathology support was provided in part by the Nancy and Buster Alvord Endowment to C.D.K. The content is solely the responsibility of the authors and does not necessarily represent the official views of NIH and its subsidiary institutes. This work was funded by the Allen Institute for Brain Science. We dedicate this paper to the vision, encouragement, and long-term support of our founder, Paul G. Allen.

## Extended Data Figures

**Figure S1:**
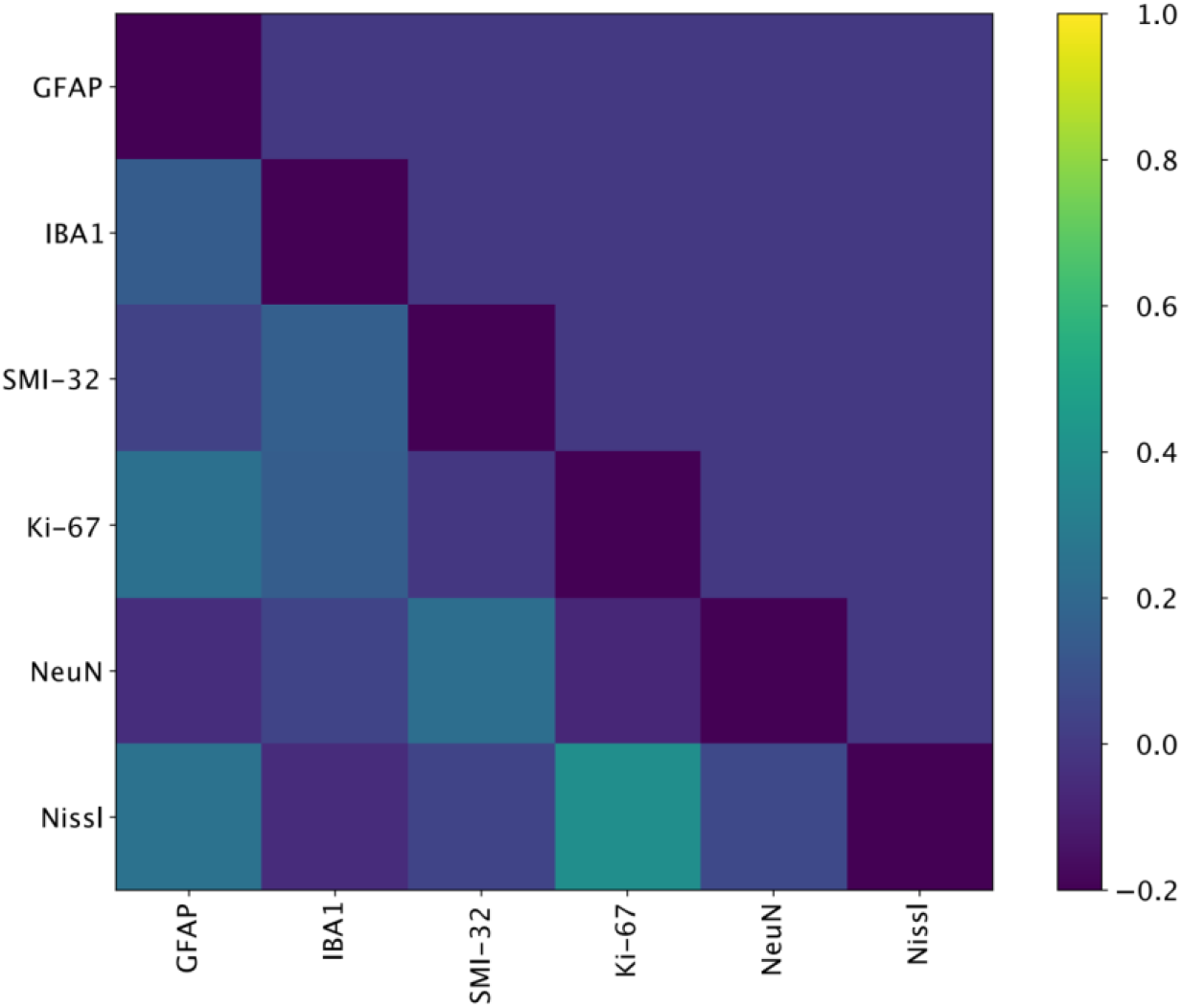
Correlation between the pathology scores: Pearson correlation coefficient between various pathology scores: GFAP, IBA1, SMI-32, Ki-67, NeuN and Nissl.

**Figure S2:**
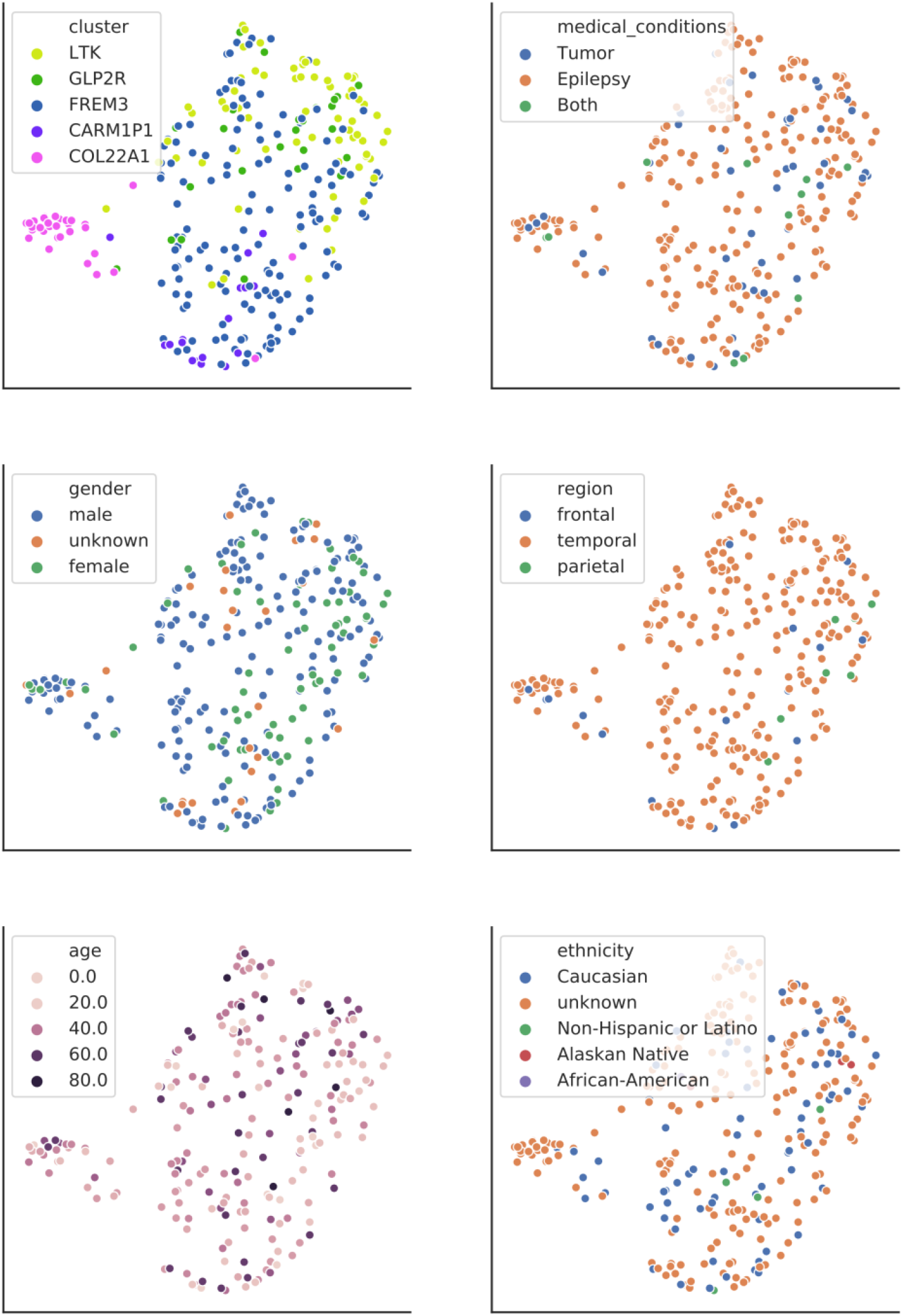
Effects of patient metadata on electrophysiology. UMAP projection of 18 electrophysiological features, with data points for each neuron colored by t-type (upper left) and by patient characteristics. In particular, cells split by medical condition (upper right) show a lack of correspondence between pathology and electrophysiology.

**Figure S3:**
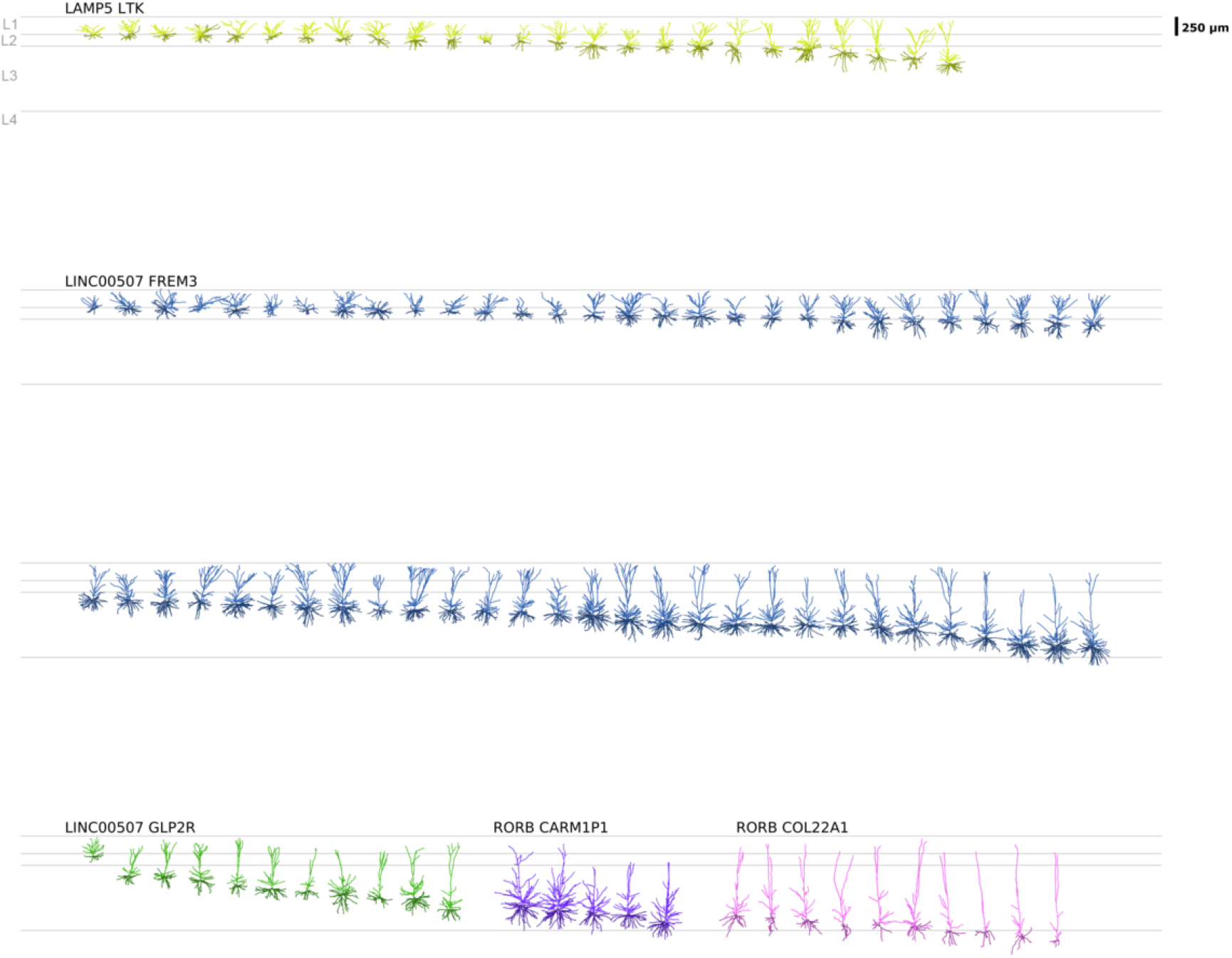
Human L2 and L3 excitatory neuron dendritic reconstructions. All human L2 and L3 excitatory neuron dendritic reconstructions ordered by t-type and aligned by layer to an average cortical template. Apical dendrites are in darker colors, basal dendrites in lighter colors.

**Figure S4:**
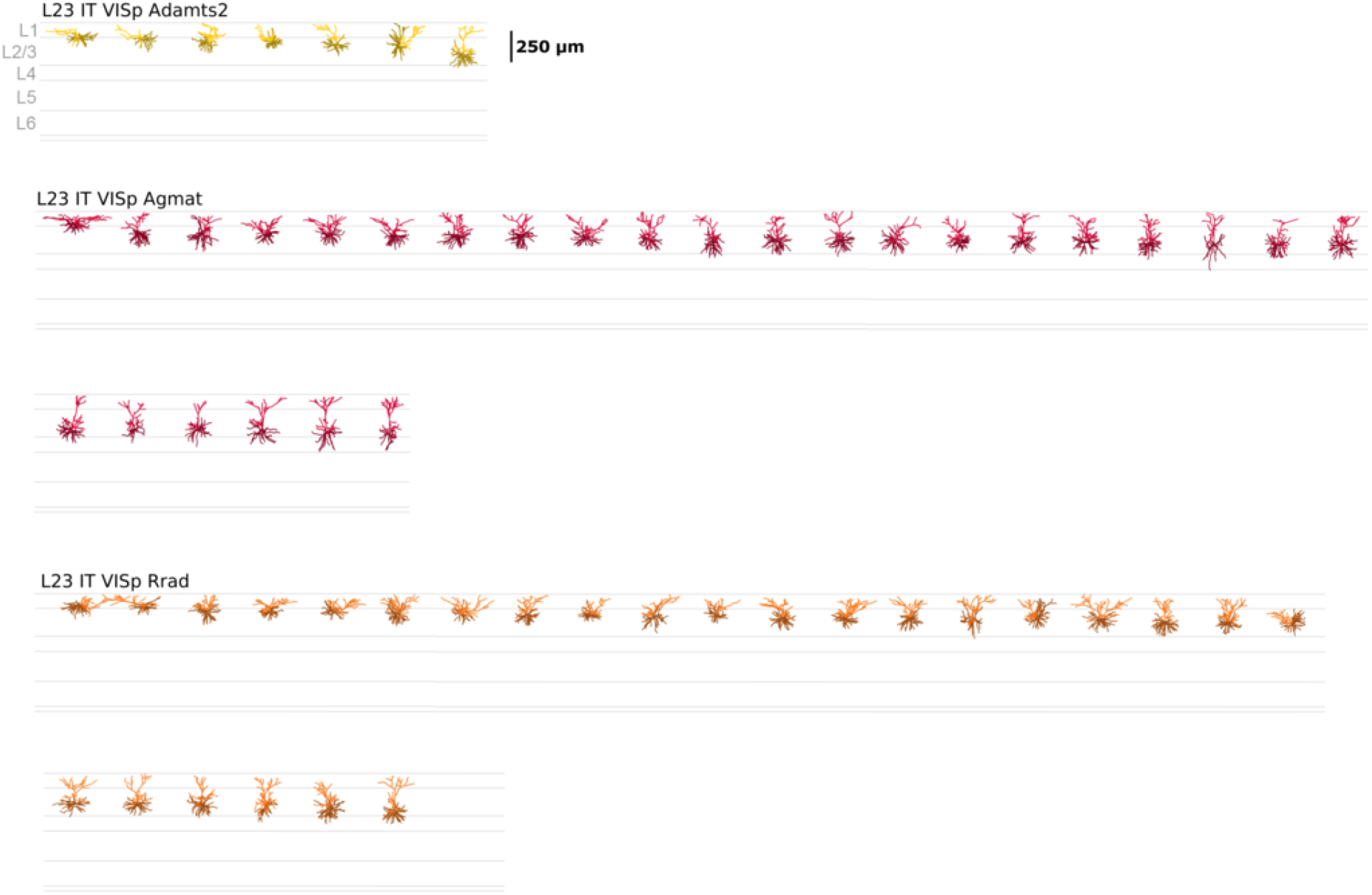
Mouse L2/3 excitatory neuron dendritic reconstructions. All mouse L2/3 excitatory neuron dendritic reconstructions ordered by t-type and aligned by layer to an average cortical template. Apical dendrites are in darker colors, basal dendrites in lighter colors.

**Figure S5:**
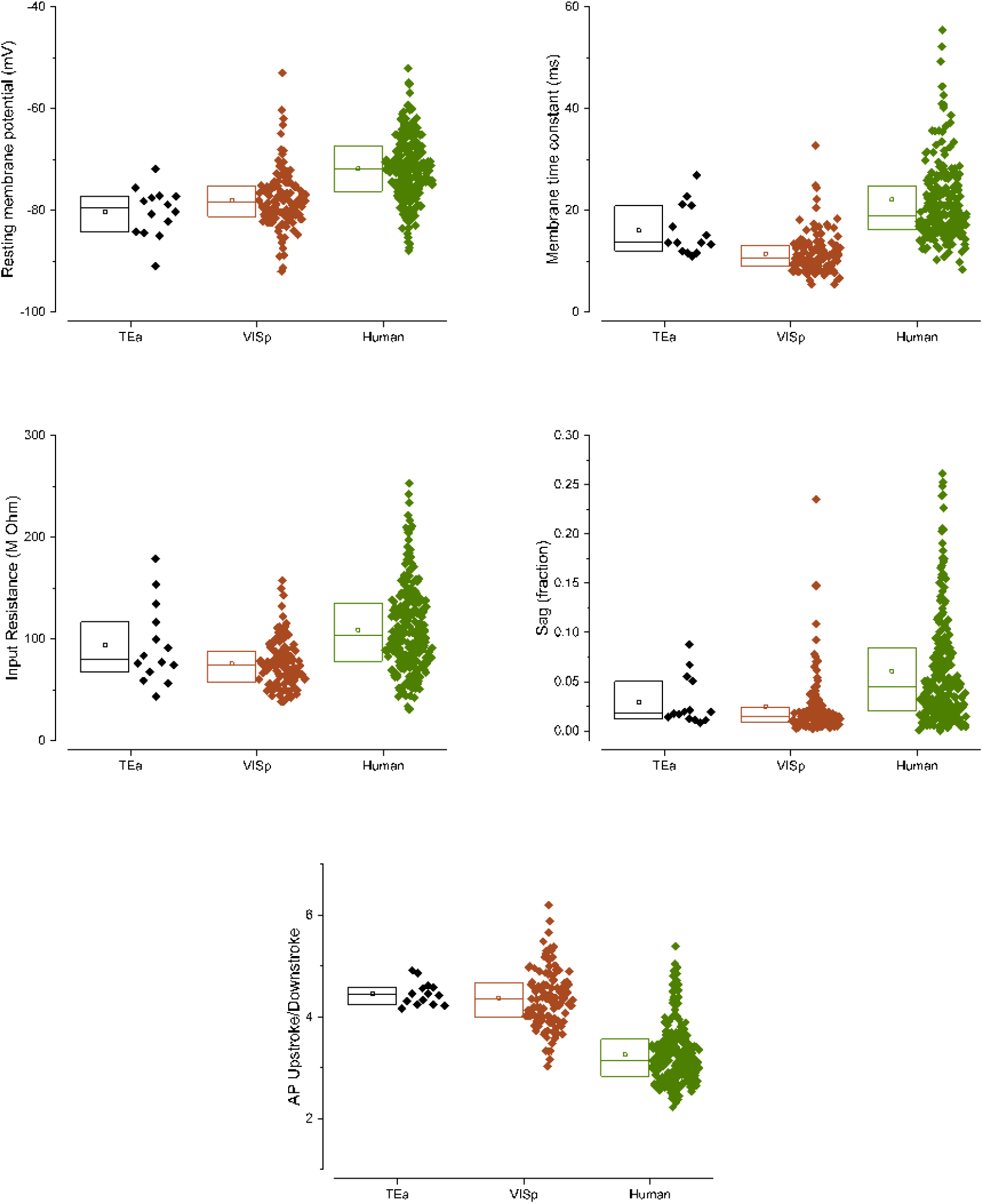
Differences in electrophysiology properties between mouse areas is smaller than those seen between mouse and human neurons. Selection of key electrophysiological features recorded from L2/3 of mouse VISp and TEa, compared to L2 and L3 of human cortex.

**Figure S6:**
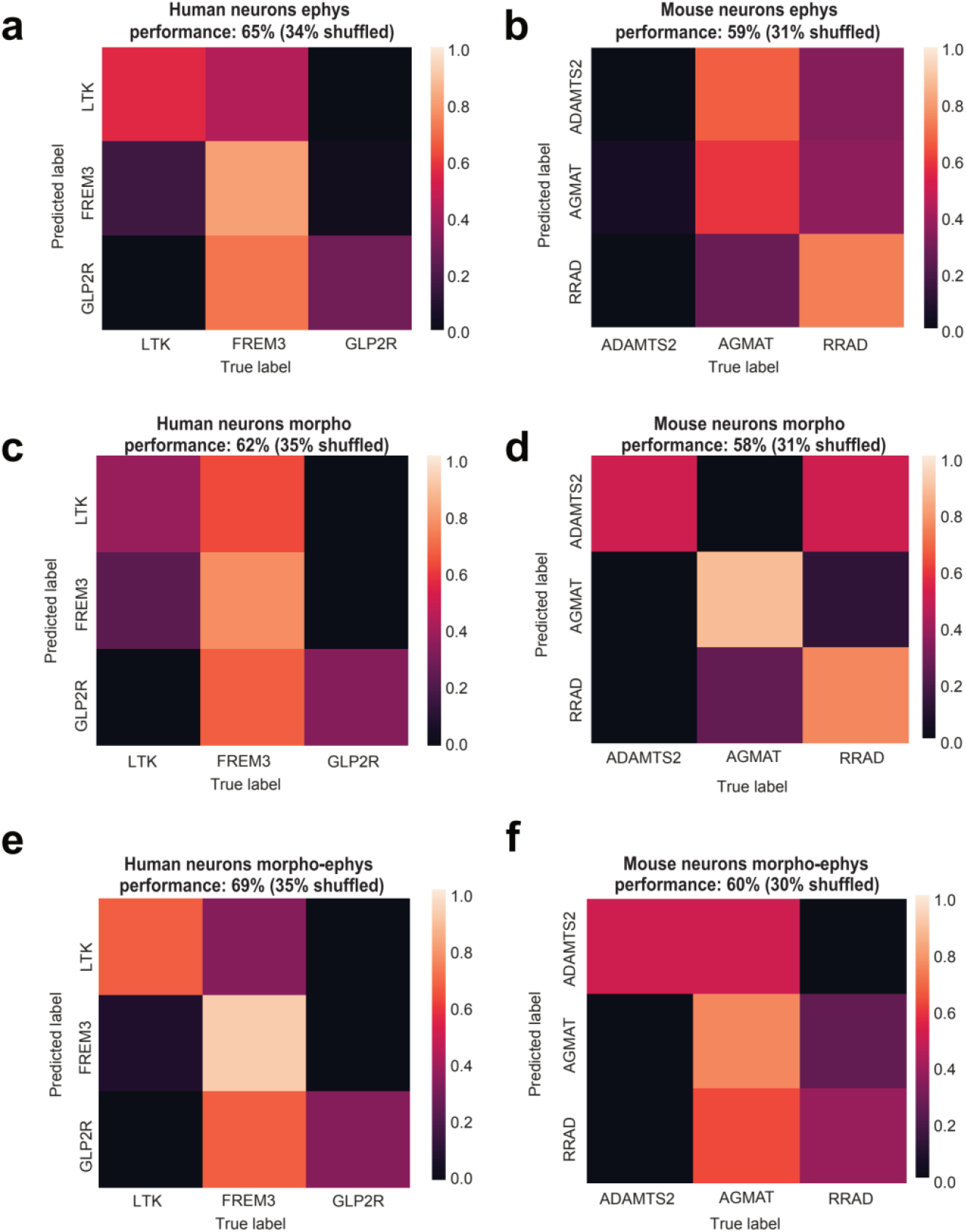
Random forest classification of t-types. **(a, b)** Confusion matrix of the random forest classifier for human and mouse neurons based on electrophysiological features. All matrices are normalized by row. **(c, d)** Confusion matrices of random forest classifiers for morphological data for mouse and human neurons. **(e, f)** Confusion matrix for the combined morpho-electric classifier. Classification performance is shown over random performance. For human ephys features the most important features were: Rin, tau, AP threshold, sag and adaptation. For mouse ephys features were: AP up/down, AP height, int. ISI, RMP and AP up. For human morpo classifier the most important features were: apical_dendrite_extent_y, basal_dendrite_extent_x_over_y, basal_dendrite_total_volume, basal_dendrite_soma_surface, basal_dendrite_emd_with_apical_dendrite. For mouse morpho classifier the most important features were: apical_dendrite_pct_intersect_basal_dendrite, apical_dendrite_early_branch, apical_dendrite_bias_x, apical_dendrite_soma_percentile_x, apical_dendrite_over_basal_dendrite_ratio_xy.

**Figure S7:**
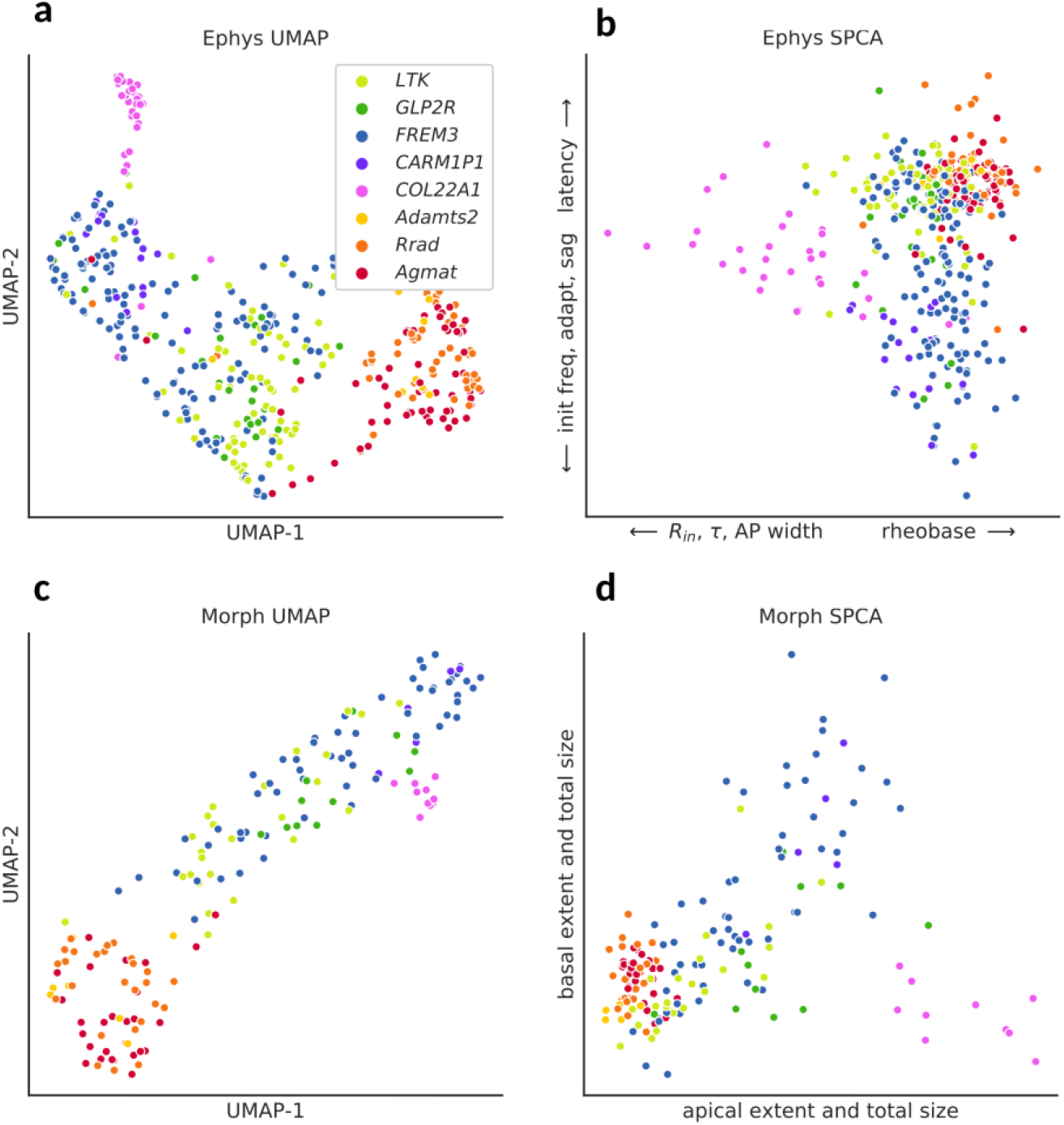
Visualization of all t-types in combined feature spaces. Projection of full electrophysiology (top) and morphology (bottom) feature spaces into two dimensions using UMAP (left) and SPCA (right).

**Figure S8:**
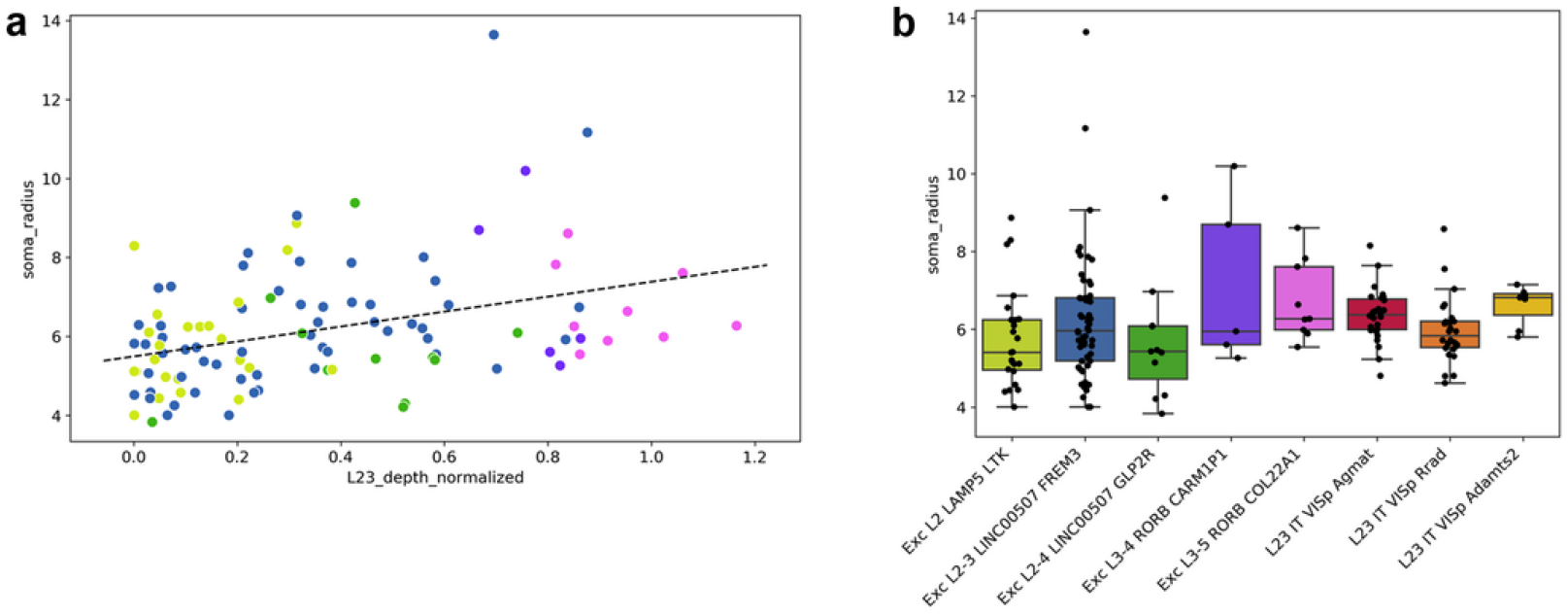
Somata radius by depth and t-type. (**a**) Soma radius vs. normalized L2,3 depth. Each soma is colored by t-type. (**b**) Average soma radius by t-type for human and mouse.

**Figure S9 (previous page):**
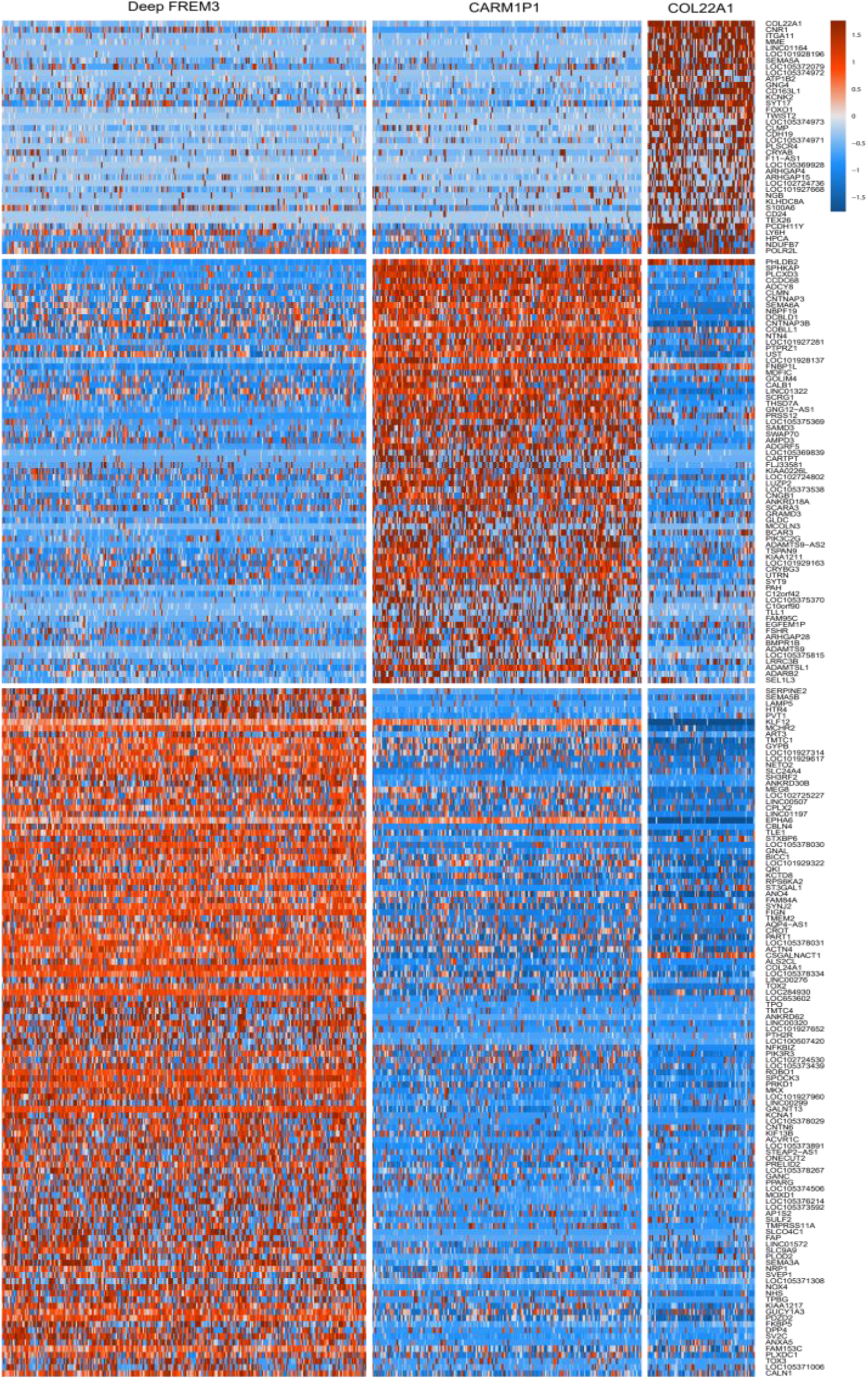
DE genes selective for one or two of the CARM1P1, COL22A1, and deep FREM3 t-types, selected using genesorteR.

**Figure S10:**
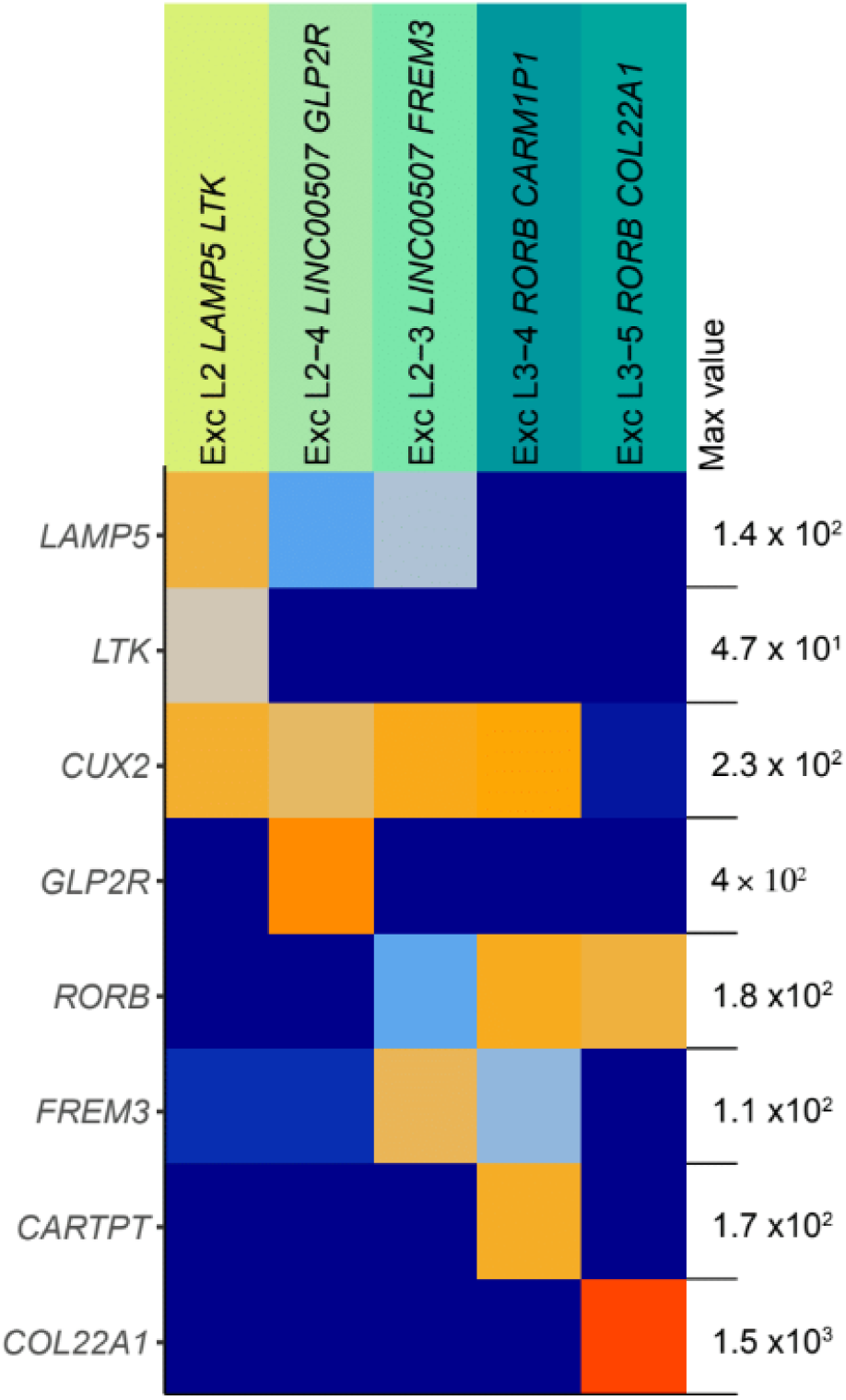
Marker gene expression shown for all five human t-types, normalized by gene.

## Extended Data Tables

**Table S1:** Cross-species comparison of electrophysiology. For homologous t-types, results are shown for unpaired t-tests across species for each feature: FDR-corrected p-values as well as Cohen’s d effect size (positive if human values are larger). 17out of 18 features are significant at FDR<0.05.

| Feature | p-value (FDR-BH) | Cohen's d (human - mouse) |
| --- | --- | --- |
| adaptation | 4.6e-15 | 0.8 |
| mean firing rate | 6.68e-09 | -0.61 |
| AP downstroke rate | 0.0 | -0.47 |
| AP fast AHP | 6.39e-35 | -1.87 |
| f-I slope | 1.2e-05 | -0.46 |
| initial firing rate | 3.5e-05 | 0.75 |
| input resistance | 1.86e-17 | 1.05 |
| latency | 3.5e-05 | -0.43 |
| AP height | 0.0 | 0.26 |
| rheobase | 1.53e-09 | -0.65 |
| sag | 9.67e-14 | 0.8 |
| time constant | 5.75e-35 | 1.62 |
| AP threshold | 4.08e-05 | -0.46 |
| AP trough | 0.0 | 0.38 |
| AP upstroke/downstroke ratio | 3.23e-31 | -1.81 |
| AP upstroke rate | 1.27e-19 | -1.18 |
| resting membrane potential | 9.11e-16 | 0.97 |
| AP width | 0.08 | 0.11 |

**Table S2:**
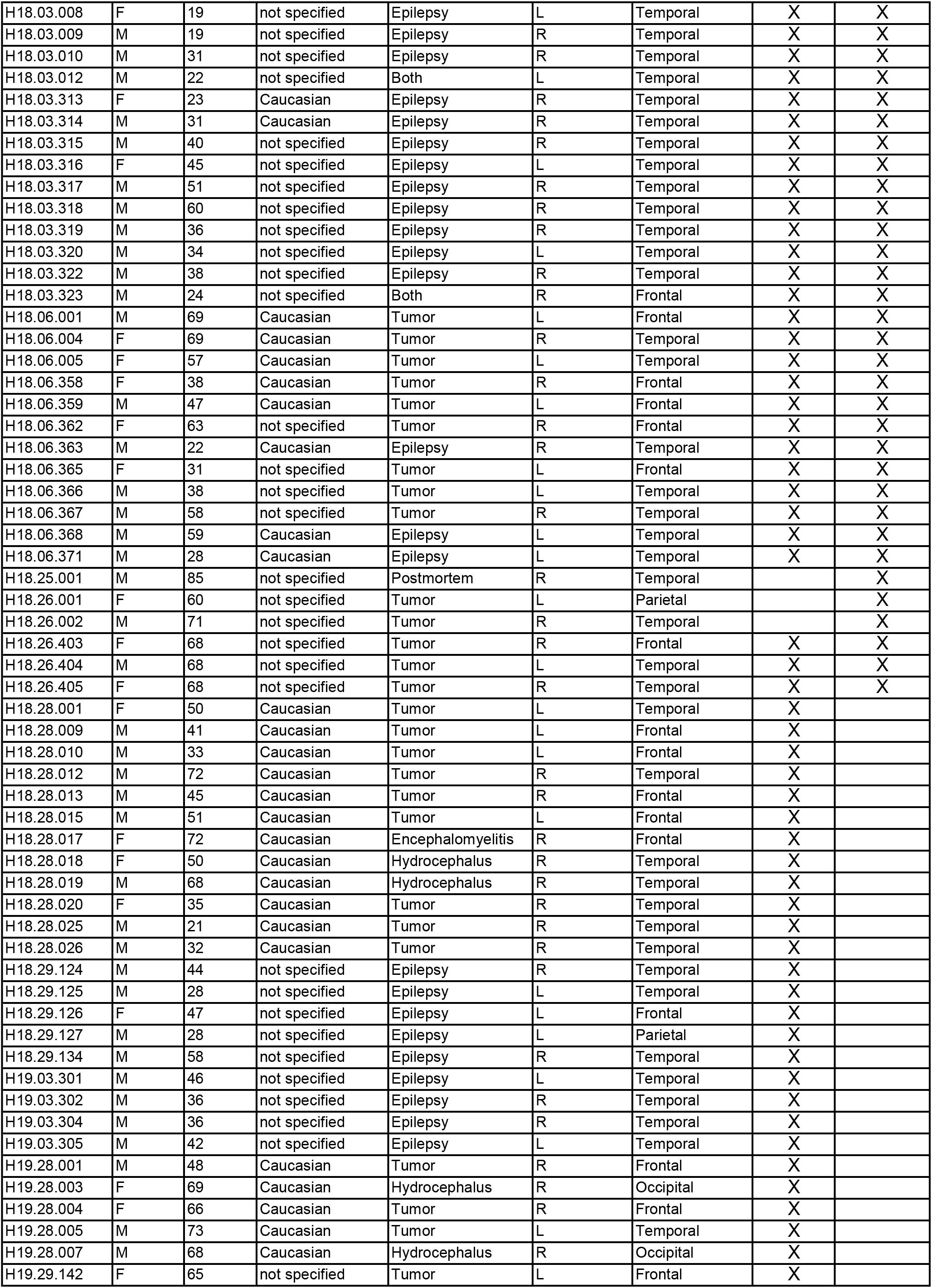
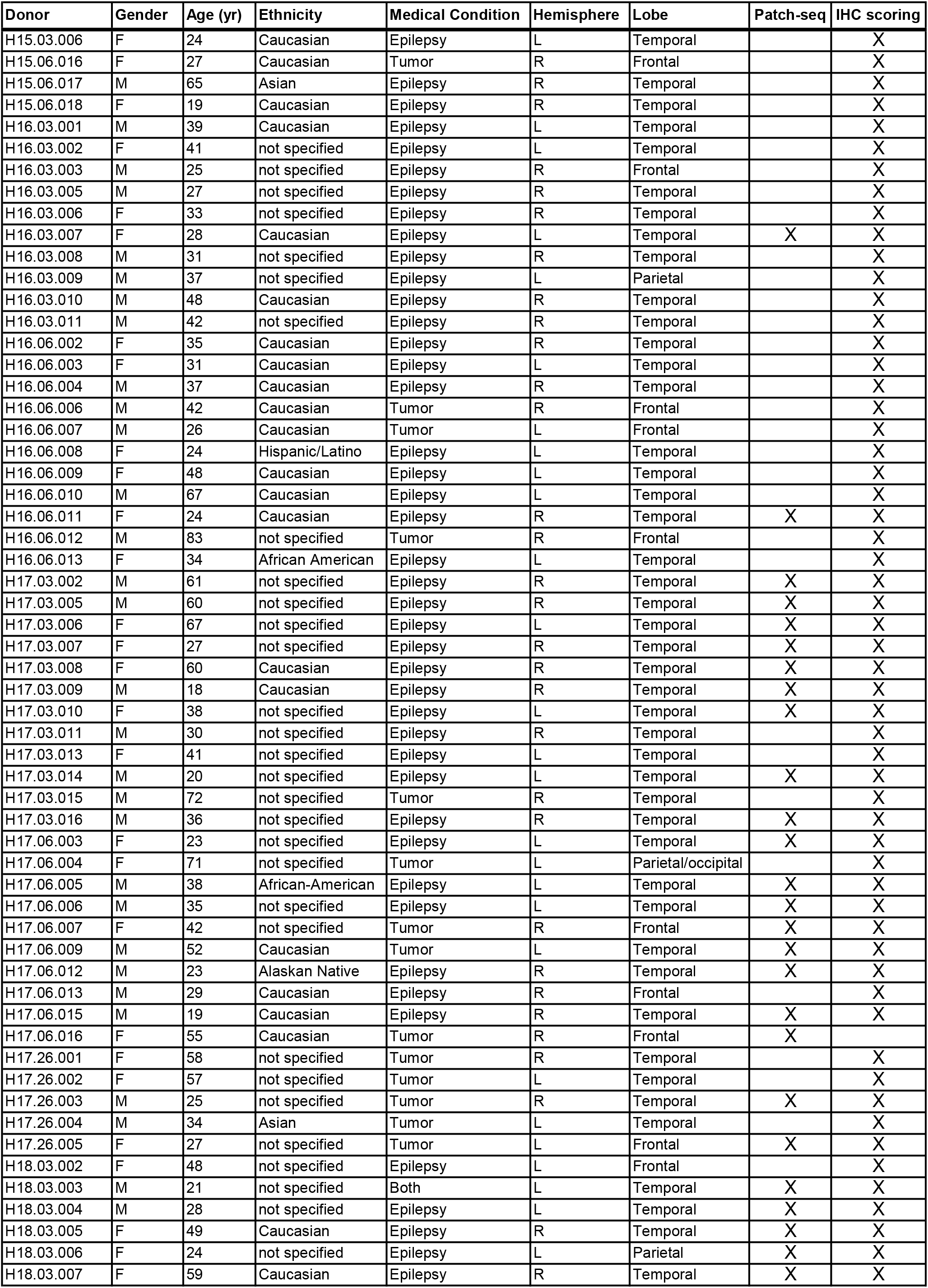
Donor attributes.

| Donor | Gender | Age (yr) | Ethnicity | Medical Condition | Hemisphere | Lobe | Patch-seq | IHC scoring |
| --- | --- | --- | --- | --- | --- | --- | --- | --- |
| H15.03.006 | F | 24 | Caucasian | Epilepsy | L | Temporal |  | X |
| H15.06.016 | F | 27 | Caucasian | Tumor | R | Frontal |  | X |
| H15.06.017 | M | 65 | Asian | Epilepsy | R | Temporal |  | X |
| H15.06.018 | F | 19 | Caucasian | Epilepsy | R | Temporal |  | X |
| H16.03.001 | M | 39 | Caucasian | Epilepsy | L | Temporal |  | X |
| H16.03.002 | F | 41 | not specified | Epilepsy | L | Temporal |  | X |
| H16.03.003 | M | 25 | not specified | Epilepsy | R | Frontal |  | X |
| H16.03.005 | M | 27 | not specified | Epilepsy | R | Temporal |  | X |
| H16.03.006 | F | 33 | not specified | Epilepsy | R | Temporal |  | X |
| H16.03.007 | F | 28 | Caucasian | Epilepsy | L | Temporal | X | X |
| H16.03.008 | M | 31 | not specified | Epilepsy | R | Temporal |  | X |
| H16.03.009 | M | 37 | not specified | Epilepsy | L | Parietal |  | X |
| H16.03.010 | M | 48 | Caucasian | Epilepsy | R | Temporal |  | X |
| H16.03.011 | M | 42 | not specified | Epilepsy | R | Temporal |  | X |
| H16.06.002 | F | 35 | Caucasian | Epilepsy | R | Temporal |  | X |
| H16.06.003 | F | 31 | Caucasian | Epilepsy | L | Temporal |  | X |
| H16.06.004 | M | 37 | Caucasian | Epilepsy | R | Temporal |  | X |
| H16.06.006 | M | 42 | Caucasian | Tumor | R | Frontal |  | X |
| H16.06.007 | M | 26 | Caucasian | Tumor | L | Frontal |  | X |
| H16.06.008 | F | 24 | Hispanic/Latino | Epilepsy | L | Temporal |  | X |
| H16.06.009 | F | 48 | Caucasian | Epilepsy | L | Temporal |  | X |
| H16.06.010 | M | 67 | Caucasian | Epilepsy | L | Temporal |  | X |
| H16.06.011 | F | 24 | Caucasian | Epilepsy | R | Temporal | X | X |
| H16.06.012 | M | 83 | not specified | Tumor | R | Frontal |  | X |
| H16.06.013 | F | 34 | African American | Epilepsy | L | Temporal |  | X |
| H17.03.002 | M | 61 | not specified | Epilepsy | R | Temporal | X | X |
| H17.03.005 | M | 60 | not specified | Epilepsy | R | Temporal | X | X |
| H17.03.006 | F | 67 | not specified | Epilepsy | L | Temporal | X | X |
| H17.03.007 | F | 27 | not specified | Epilepsy | R | Temporal | X | X |
| H17.03.008 | F | 60 | Caucasian | Epilepsy | R | Temporal | X | X |
| H17.03.009 | M | 18 | Caucasian | Epilepsy | R | Temporal | X | X |
| H17.03.010 | F | 38 | not specified | Epilepsy | L | Temporal | X | X |
| H17.03.011 | M | 30 | not specified | Epilepsy | R | Temporal |  | X |
| H17.03.013 | F | 41 | not specified | Epilepsy | L | Temporal |  | X |
| H17.03.014 | M | 20 | not specified | Epilepsy | L | Temporal | X | X |
| H17.03.015 | M | 72 | not specified | Tumor | R | Temporal |  | X |
| H17.03.016 | M | 36 | not specified | Epilepsy | R | Temporal | X | X |
| H17.06.003 | F | 23 | not specified | Epilepsy | L | Temporal | X | X |
| H17.06.004 | F | 71 | not specified | Tumor | L | Parietal/occipital |  | X |
| H17.06.005 | M | 38 | African-American | Epilepsy | L | Temporal | X | X |
| H17.06.006 | M | 35 | not specified | Epilepsy | L | Temporal | X | X |
| H17.06.007 | F | 42 | not specified | Tumor | R | Frontal | X | X |
| H17.06.009 | M | 52 | Caucasian | Tumor | L | Temporal | X | X |
| H17.06.012 | M | 23 | Alaskan Native | Epilepsy | R | Temporal | X | X |
| H17.06.013 | M | 29 | Caucasian | Epilepsy | R | Frontal |  | X |
| H17.06.015 | M | 19 | Caucasian | Epilepsy | R | Temporal | X | X |
| H17.06.016 | F | 55 | Caucasian | Tumor | R | Frontal | X |  |
| H17.26.001 | F | 58 | not specified | Tumor | R | Temporal |  | X |
| H17.26.002 | F | 57 | not specified | Tumor | L | Temporal |  | X |
| H17.26.003 | M | 25 | not specified | Tumor | R | Temporal | X | X |
| H17.26.004 | M | 34 | Asian | Tumor | L | Temporal |  | X |
| H17.26.005 | F | 27 | not specified | Tumor | L | Frontal | X | X |
| H18.03.002 | F | 48 | not specified | Epilepsy | L | Frontal |  | X |
| H18.03.003 | M | 21 | not specified | Both | L | Temporal | X | X |
| H18.03.004 | M | 28 | not specified | Epilepsy | L | Temporal | X | X |
| H18.03.005 | F | 49 | Caucasian | Epilepsy | R | Temporal | X | X |
| H18.03.006 | F | 24 | not specified | Epilepsy | L | Parietal | X | X |
| H18.03.007 | F | 59 | Caucasian | Epilepsy | R | Temporal | X | X |
| H18.03.008 | F | 19 | not specified | Epilepsy | L | Temporal | X | X |
| H18.03.009 | M | 19 | not specified | Epilepsy | R | Temporal | X | X |
| H18.03.010 | M | 31 | not specified | Epilepsy | R | Temporal | X | X |
| H18.03.012 | M | 22 | not specified | Both | L | Temporal | X | X |
| H18.03.313 | F | 23 | Caucasian | Epilepsy | R | Temporal | X | X |
| H18.03.314 | M | 31 | Caucasian | Epilepsy | R | Temporal | X | X |
| H18.03.315 | M | 40 | not specified | Epilepsy | R | Temporal | X | X |
| H18.03.316 | F | 45 | not specified | Epilepsy | L | Temporal | X | X |
| H18.03.317 | M | 51 | not specified | Epilepsy | R | Temporal | X | X |
| H18.03.318 | M | 60 | not specified | Epilepsy | R | Temporal | X | X |
| H18.03.319 | M | 36 | not specified | Epilepsy | R | Temporal | X | X |
| H18.03.320 | M | 34 | not specified | Epilepsy | L | Temporal | X | X |
| H18.03.322 | M | 38 | not specified | Epilepsy | R | Temporal | X | X |
| H18.03.323 | M | 24 | not specified | Both | R | Frontal | X | X |
| H18.06.001 | M | 69 | Caucasian | Tumor | L | Frontal | X | X |
| H18.06.004 | F | 69 | Caucasian | Tumor | R | Temporal | X | X |
| H18.06.005 | F | 57 | Caucasian | Tumor | L | Temporal | X | X |
| H18.06.358 | F | 38 | Caucasian | Tumor | R | Frontal | X | X |
| H18.06.359 | M | 47 | Caucasian | Tumor | L | Frontal | X | X |
| H18.06.362 | F | 63 | not specified | Tumor | R | Frontal | X | X |
| H18.06.363 | M | 22 | Caucasian | Epilepsy | R | Temporal | X | X |
| H18.06.365 | F | 31 | not specified | Tumor | L | Frontal | X | X |
| H18.06.366 | M | 38 | not specified | Tumor | L | Temporal | X | X |
| H18.06.367 | M | 58 | not specified | Tumor | R | Temporal | X | X |
| H18.06.368 | M | 59 | Caucasian | Epilepsy | L | Temporal | X | X |
| H18.06.371 | M | 28 | Caucasian | Epilepsy | L | Temporal | X | X |
| H18.25.001 | M | 85 | not specified | Postmortem | R | Temporal |  | X |
| H18.26.001 | F | 60 | not specified | Tumor | L | Parietal |  | X |
| H18.26.002 | M | 71 | not specified | Tumor | R | Temporal |  | X |
| H18.26.403 | F | 68 | not specified | Tumor | R | Frontal | X | X |
| H18.26.404 | M | 68 | not specified | Tumor | L | Temporal | X | X |
| H18.26.405 | F | 68 | not specified | Tumor | R | Temporal | X | X |
| H18.28.001 | F | 50 | Caucasian | Tumor | L | Temporal | X |  |
| H18.28.009 | M | 41 | Caucasian | Tumor | L | Frontal | X |  |
| H18.28.010 | M | 33 | Caucasian | Tumor | L | Frontal | X |  |
| H18.28.012 | M | 72 | Caucasian | Tumor | R | Temporal | X |  |
| H18.28.013 | M | 45 | Caucasian | Tumor | R | Frontal | X |  |
| H18.28.015 | M | 51 | Caucasian | Tumor | L | Frontal | X |  |
| H18.28.017 | F | 72 | Caucasian | Encephalomyelitis | R | Frontal | X |  |
| H18.28.018 | F | 50 | Caucasian | Hydrocephalus | R | Temporal | X |  |
| H18.28.019 | M | 68 | Caucasian | Hydrocephalus | R | Temporal | X |  |
| H18.28.020 | F | 35 | Caucasian | Tumor | R | Temporal | X |  |
| H18.28.025 | M | 21 | Caucasian | Tumor | R | Temporal | X |  |
| H18.28.026 | M | 32 | Caucasian | Tumor | R | Temporal | X |  |
| H18.29.124 | M | 44 | not specified | Epilepsy | R | Temporal | X |  |
| H18.29.125 | M | 28 | not specified | Epilepsy | L | Temporal | X |  |
| H18.29.126 | F | 47 | not specified | Epilepsy | L | Frontal | X |  |
| H18.29.127 | M | 28 | not specified | Epilepsy | L | Parietal | X |  |
| H18.29.134 | M | 58 | not specified | Epilepsy | R | Temporal | X |  |
| H19.03.301 | M | 46 | not specified | Epilepsy | L | Temporal | X |  |
| H19.03.302 | M | 36 | not specified | Epilepsy | R | Temporal | X |  |
| H19.03.304 | M | 36 | not specified | Epilepsy | R | Temporal | X |  |
| H19.03.305 | M | 42 | not specified | Epilepsy | L | Temporal | X |  |
| H19.28.001 | M | 48 | Caucasian | Tumor | R | Frontal | X |  |
| H19.28.003 | F | 69 | Caucasian | Hydrocephalus | R | Occipital | X |  |
| H19.28.004 | F | 66 | Caucasian | Tumor | R | Frontal | X |  |
| H19.28.005 | M | 73 | Caucasian | Tumor | L | Temporal | X |  |
| H19.28.007 | M | 68 | Caucasian | Hydrocephalus | R | Occipital | X |  |
| H19.29.142 | F | 65 | not specified | Tumor | L | Frontal | X |  |

